# Identification of Novel DNA Sequence Motifs that Modulate Transcription in T cells

**DOI:** 10.1101/2025.07.08.660448

**Authors:** Nicole Knoetze, Eric Yung, Anthony Bayega, Scott D. Brown, Robert A. Holt

## Abstract

Considerable progress has been made towards associating transcription factor binding sites (TFBS) with cell-type-specific gene expression, however, the full repertoire of DNA sequence motifs that regulate transcription remains unknown. Improving our understanding of transcriptional regulation is especially important in T cells, given the enormous potential of genetically engineered T cells as an emerging class of therapeutics. Here, we report results from a comprehensive and unbiased survey investigating whether there are novel motifs enriched in regulatory regions of genes with the highest constitutive and selective expression across diverse T-cell subsets. Using computational and experimental methods, we identified 2,036 novel motifs and 629 previously curated TFBS that are enriched, both individually and in specific combinations, in the regulatory regions of genes exhibiting T-cell-specific gene expression. We then used the self-transcribing active regulatory region sequencing (STARR-seq) assay to evaluate all possible three-way combinations of a subset of 18 candidate motifs to test their ability to modulate transcription in immortalized lymphoblastic cell lines of T-cell origin (Jurkat E6) versus myeloid origin (K562). Our results revealed novel motifs that modulate gene transcription in T cells, with some exhibiting stronger regulatory effects than TFBS for TFs with established roles in T cells. The regulatory activity of these novel motifs was influenced by the motif’s orientation, position, and copy number. Overall, these results highlight our incomplete understanding of the relationship between sequence composition and T-cell gene regulation and indicate that previously annotated TFBS represent only a subset of motifs capable of modulating gene transcription in T cells.

## Introduction

The precise control of cell-type-specific gene expression is thought to be dictated by the combinatorial action of transcription factors (TFs) bound to distinct DNA sequence motifs (Transcription Factor Binding Sites, TFBS) within open chromatin regions^1–4^. Identifying the exact DNA sequence elements capable of regulating gene expression in a cell-type-specific manner is essential for advancing our knowledge of gene regulation and for designing synthetic regulatory sequences for gene therapy. While many studies have identified TFBS associated with cell-type-specific gene expression, there has been limited emphasis on comprehensively assessing sequence composition for regulatory potential, and no study has done this exhaustively. Further, although many studies have performed *de novo* motif discovery and have identified novel motifs within putative regulatory regions, the functional relevance of these motifs typically goes untested^2,5,6^. This gap is particularly notable in T cells, where, despite their central importance to immunotherapy applications, the regulatory landscape governing T-cell-specific gene expression remains largely unexplored, particularly with respect to novel motifs. Thus, we asked if there are previously unaccounted for sequence motifs that are associated with the transcriptional control of genes with the highest level of cell-type-specific expression, and if so, how prevalent are they, and do they adhere to a regulatory grammar defined by order, orientation, and copy number^4,7^.

To pursue this line of enquiry, we developed a computational and experimental framework comprising four main parts: i) identifying genes with the highest, constitutive and selective expression across various subsets of non-activated T cells, ii) determining the putative regulatory regions of these genes, iii) performing a comprehensive and unbiased search for enriched motifs within these regulatory regions, and iv) performing a deep functional screening of candidate motifs and combinations of motifs in cell lines using a massively parallel reporter assay (STARR-seq^8^). Using this approach, we identified combinations of novel motifs and previously curated TFBS that are significantly enriched in the regulatory regions of genes exhibiting T-cell-specific gene expression. From the STARR-seq assay, we identified several novel motifs that were able to modulate gene transcription in T cells, in some instances at levels higher than those observed for curated binding sites for TFs with well-established roles in T cells. Moreover, we observed a regulatory grammar for these motifs, similar to that for TFBS^7,9,10^ highlighting preferences for order, orientation, and copy number. Overall, this work highlights the utility of our approach, which can be applied to any cell type, and underscores the importance of expanding motif discovery to include novel motifs to better understand cell-type-specific gene expression. Despite these advances, the mechanisms by which the identified novel motifs alter transcription in T cells, and whether this may involve specific trans-acting factors, remain unknown and will require further study.

## Results

### Identifying Genes with T-cell-specific Expression and Their Associated Regulatory Regions

T cells encompass a heterogeneous population of immune-cell subtypes defined by the expression of specific marker genes^11^. To identify sequence features that may be potentially relevant to the regulation of T-cell-specific genes, we interrogated a published RNA-seq dataset from the DICE consortium^12^ comprising nine T-cell subtypes (naive CD4^+^ T cells, naive CD8^+^ T cells, naive regulatory T cells, memory regulatory T cells, Th17, Th1-17, Th1, Th2, and follicular helper T cells) from 91 healthy individuals. We included a variety of T-cell subsets to increase the likelihood that results would be relevant to T cells as a whole and not just a particular subset. For comparisons to cell types other than T cells, we utilized additional RNA-seq datasets for other immune cells, reproductive cells, hepatocytes, hematopoietic stem cells, and endothelial cells (**Supplemental Figure S1**). Overall, our comprehensive dataset included 1,740 RNA-seq samples spanning 22 cell types (**Supplemental Table S1**).

Using these data resources, we first identified the genes with the highest constitutive and selective expression levels across T-cell subtypes. Briefly, we summed the expression values across all T-cell and non-T-cell cell samples separately to obtain the aggregate expression value for each protein-coding gene in each of these two sample sets. From the aggregate expression values in T cells, we determined the median aggregate value (aggregate expression value = 237.04) and removed genes below this value. Next, we ranked the remaining genes (n = 10,104) based on their aggregate expression values from high to low in T cells and from low to high in non-T-cell samples. We then summed these two ranks to calculate a T-cell specificity score for each gene representing its level of enrichment in T cells, where low values represent genes that are highly enriched in T cells. For subsequent analysis, we further partitioned the genes into specific gene sets by setting thresholds based on the mean and standard deviation (SD) of the distribution of specificity scores, where a low specificity score indicates greater T-cell specificity. To create a zone of exclusion between our gene sets, we established two thresholds. The first threshold was set at four SDs below the mean, and we classified genes with specificity scores below this threshold as our top-ranked T-cell genes (n = 22; top-ranked, **Supplemental Figure S2A**). The second threshold was set at three SDs below the mean, where we defined genes with specificity scores above this threshold as a part of the comparator gene set (n = 8,562; comparator set, **Supplemental Figure S2A**). Finally, we defined a size-matched control gene set comprising the 22 bottom-ranked genes (bottom-ranked, **Supplemental Figure S2B**). In characterizing these gene sets, we noted that the 22 top-ranked genes exhibited high, consistent and selective gene expression across multiple T-cell subtypes, while the 22 bottom-ranked genes demonstrated a heterogeneous gene expression profile (**Figure 1A**). On average, the top-ranked genes had an expression level of 2,654 (normalised counts; DESeq, see Methods) in T cells compared to 12.2 in non-T cells, while the 22 bottom-ranked genes had an expression level of 1,519 in non-T cells and 47.3 in T cells. Gene ontology term analysis indicated that the 22 top-ranked genes were associated with biological processes characteristic of T cells, including T-cell activation, differentiation, and T-cell receptor signalling (**Figure 1B**, Benjamini-Hochberg adjusted, p-value < 0.05). Conversely, the 22 bottom-ranked genes were associated with immune-related biological processes, but not T-cell biological processes specifically (**Supplemental Figure S3**). In summary, our methodology clearly discriminated sets of genes with and without T-cell-specific expression.

**Figure 1.**
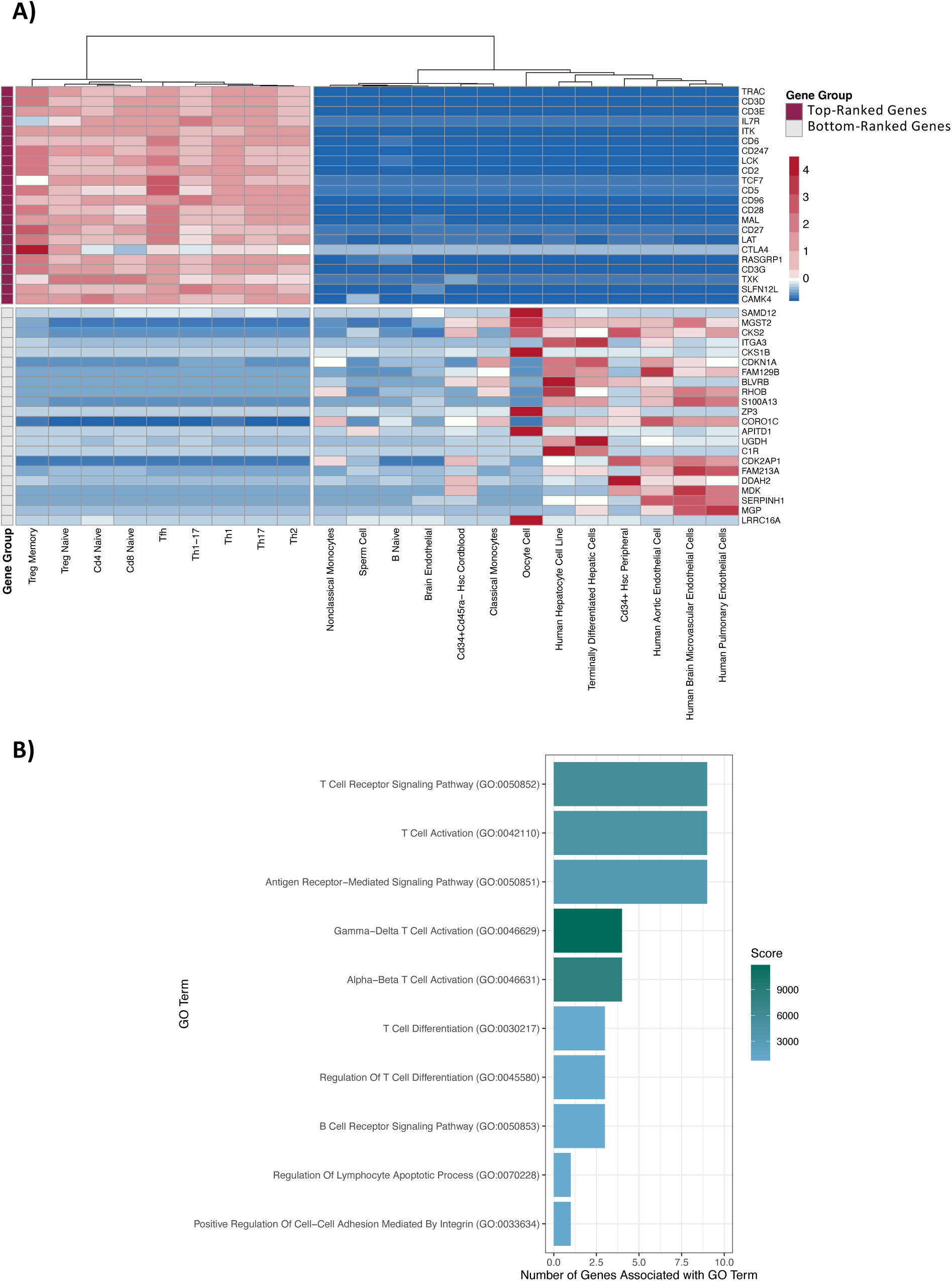
Expression of sets of ranked genes in T cells. A) The median expression levels for the 22 top-ranked and the 22 bottom-ranked genes. Genes were given a specificity score that is based on having high expression in T cells and low expression in non-T cells, and ranked, as described in the Results. This plot shows the 22 top-ranked genes and the 22 bottom-ranked T-cell genes in rows, and their corresponding expression profiles across various cell types in columns. B) The 22 top-ranked genes are associated with T-cell-related biological processes. The top ten enriched GO terms are shown (y-axis, Benjamini-Hochberg adjusted p-value < 0.05). The bar plot indicates how many of the top-ranked genes (x-axis) are linked to each GO term, with the colour corresponding to the score derived from the GO term analysis. GO terms are arranged in order of their adjusted p-value.

Active regulatory DNA typically coincides with open chromatin regions, which are hypersensitive to DNase1 digestion. Therefore, to identify putative regulatory regions for T-cell-expressed genes we interrogated a published CD4^+^ and CD8^+^ T cell DNase-seq dataset^13^ (n = 134,233 regulatory regions). After restricting the DNase-seq defined regulatory regions to only those annotated for both cell types (CD4^+^ and CD8^+^), we retained 130,838 regulatory regions (97.5%), averaging 240 bp in length (range: 42 - 2,428 bp, standard deviation: 84 bp). We then classified these regions as proximal or distal, which are the two main categories of regulatory sequences^14,15^. Proximal regions were arbitrarily defined as being within 500 bp of a protein-coding gene’s transcription start site. Unlike proximal regulatory regions, distal regulatory regions are more difficult to define as they may be located thousands of kilobases, or further, from the genes they regulate^16^ and are brought into the proximity of their target genes through DNA looping^17,18^. Fortuitously, however, chromosome conformation assays such as Hi-C and Hi-ChIP that capture these dynamics have greatly improved our ability to identify the target genes of distal regulatory regions^17,18^. Therefore, to link distal regulatory regions to their target genes, we utilized an H3K27ac Hi-ChIP dataset^19^ for CD8^+^ and CD4^+^ T cells. Briefly, for each gene, we obtained all annotated genomic interactions from the Hi-ChIP dataset containing the gene’s proximal regulatory regions. Distal regulatory regions contained within the same interaction annotation were assigned as the distal regulatory regions for the respective gene (see Methods). On average, the 22 top-ranked genes were associated with 1.95 +/-0.99 (mean+/- SD) proximal regulatory regions and linked to an average of 18.5+/- 11.7 (mean +/- SD) distal regulatory regions. The 22 bottom-ranked genes were associated with 2.27 +/- 1.12 proximal and 9.14 +/- 10.1 distal regulatory regions. Ultimately, integrating RNA-seq, DNase-seq, and H3K27ac Hi-ChIP datasets led to the identification of 43 proximal and 390 distal regulatory regions for the 22 top-ranked genes, and 50 proximal and 197 distal regulatory regions for the 22 bottom-ranked T-cell genes. The similarity of these values suggests that differential regulation of the two gene sets is more likely attributable to the sequence composition of their associated regulatory regions, rather than the number of associated regulatory regions.

### Novel DNA Sequence Motifs are Selectively Enriched in Regulatory Regions for the Top-Ranked T-cell-specific Genes

Having annotated putative regulatory regions, we investigated whether these regions contain novel DNA sequence motifs. To search for motifs, we decomposed the sequences of the regulatory regions for the 22 top-ranked genes into all possible 12-mers, the average size of motifs in JASPAR^20^. We then clustered similar 12-mers using a hamming distance of one and generated Positional Probability Matrices (PPMs, n=13,576), representing motifs, from the DNA sequences in each cluster. To focus on putative novel PPMs, we removed any PPMs that resembled a TFBS from the JASPAR database^20^. After clustering redundant PPMs, we had 4,934 PPMs representing novel motifs (**Figure 2A**).

**Figure 2.**
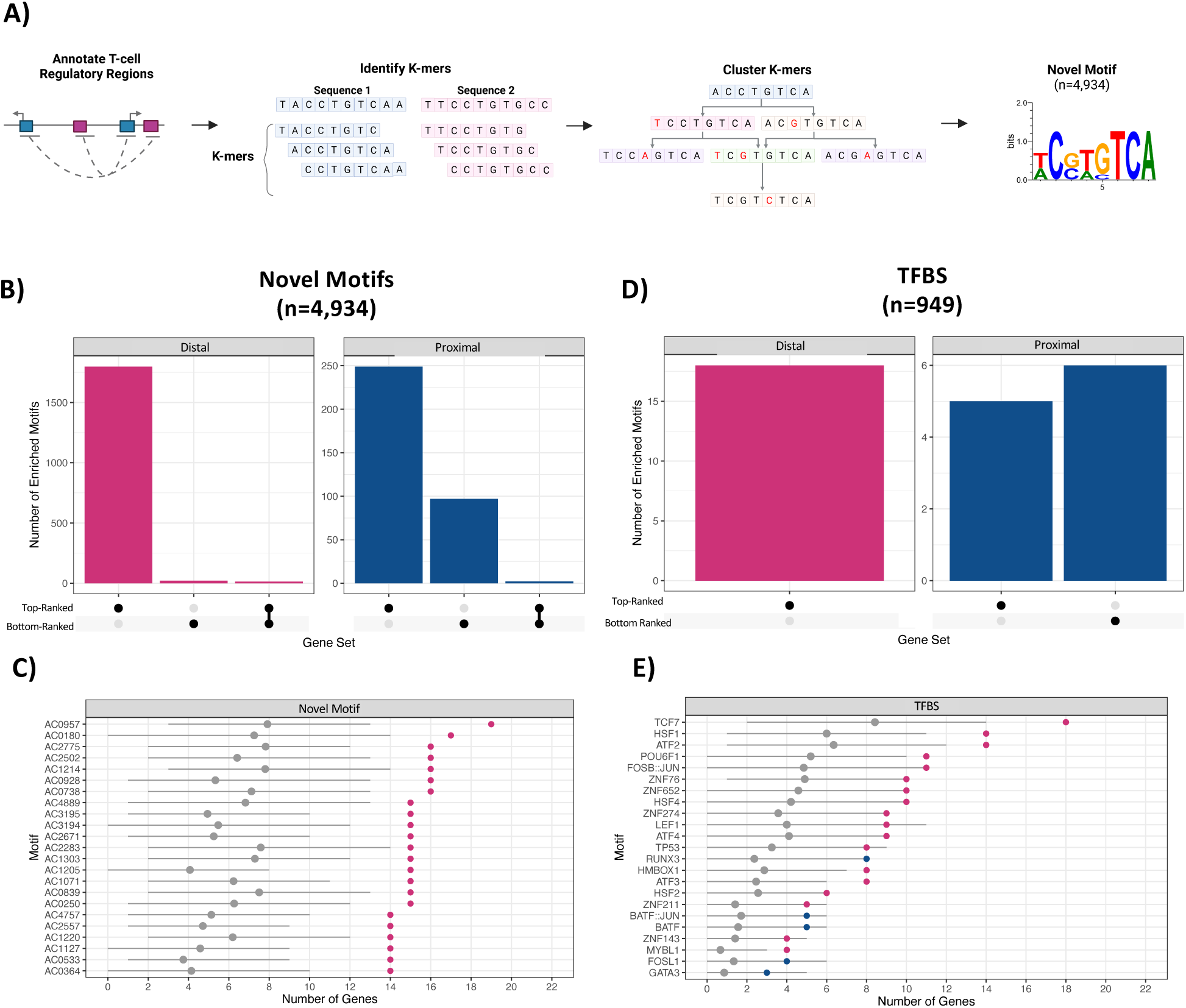
Motifs enriched in the regulatory regions of the top and bottom-ranked T-cell genes. A) Schematic representing the workflow for identifying novel motifs. B) The number of enriched novel motifs (y-axis) by regulatory region type (proximal or distal) and gene set (x-axis). C) The top 23 novel motifs enriched in regulatory regions associated with the 22 top-ranked genes (p-value < 0.05, fold change >2) are shown. Motifs are ordered by the number of top-ranked genes they are associated with. Grey points indicate the average number of genes from the comparator gene set that contain a site for the given motif (y-axis). D) The number of enriched TFBS from JASPAR (y-axis) by regulatory region type and the gene set (x-axis). E) The 23 TFBS motifs enriched in regulatory regions associated with the 22 top-ranked genes (p-value < 0.05, fold change >2) are shown (right). Motifs are ordered by the number of top-ranked genes they are associated with. Grey points indicate the average number of comparator genes containing a site for the given motif (y-axis).

To test if these novel motifs (n=4,934) are more commonly found in regulatory regions for the 22 top-ranked genes compared to the comparator set of 8,562 non-specific T-cell genes, we developed a Monté Carlo statistical framework. The framework compares the number of T-cell genes from the top-ranked gene set whose regulatory regions contain at least one instance of the motif to a distribution obtained from 10,000 random samplings of 22 non-specific T-cell genes from the comparator gene set (**Supplemental Figure S4**). Motifs that had a nominal p-value < 0.05 and were associated with twice as many top-ranked genes as iteratively sampled comparator set genes were designated as enriched (n=2,036), and all remaining motifs were designated as non-enriched (n=2,898). Of the enriched novel motifs, 120 were enriched in both distal and proximal regulatory regions. For the 22 bottom-ranked genes, we only identified 151 enriched novel motifs, 16 of which were also enriched in regulatory regions for the 22 top-ranked genes (**Figure 2B-C**). Notably, 98.9% of the enriched motifs were found in regulatory regions for the 22 top-ranked genes at a frequency higher than would be expected based on their distribution across the genome (**Supplemental Figure S5**). In contrast, 47.4% of the non-enriched motifs were found in regulatory regions for the 22 top-ranked genes at a frequency higher than would be expected. These results highlight that the enriched novel motifs are not randomly distributed across the genome and are concentrated in the regulatory regions associated with genes with T-cell-specific expression.

### Regulatory Regions for Top-Ranked T-cell-specific Genes are Enriched for Binding Sites for T-cell Transcription Factors

In a separate analysis, we used the same statistical framework to analyze known TFBS from the JASPAR database (n=949) and identified 23 TFBS motifs enriched in the regulatory regions of the 22 top-ranked genes (**Figure 2D-E**). Notably, the TFBS enriched in distal and proximal regulatory regions were distinct, suggesting that there may be different proximal and distal mechanisms of regulation (**Figure 2E**). Many of the enriched TFBS are recognized as binding sites for TFs with established roles in T-cell biology, including TCF7 and LEF1, which are key T-cell developmental TFs^21–23^. We also identified enrichment of binding sites for the FOSB::JUN dimer, an important component of the AP-1 complex, which is involved in a variety of processes in T cells including T-cell activation and proliferation^24,25^. These results highlight the ability of our approach to identify enriched motifs in regulatory regions for cell-type-specific genes. Lastly, in our control gene set containing the 22 bottom-ranked genes, only six TFBS were enriched in proximal regulatory regions (**Figure 2D**) and no TFBS were enriched in distal regulatory regions. Thus, the regulatory regions of the 22 top-ranked genes appear to share specific sequence composition, and their expression may be influenced, at least partially, by novel motifs. These novel motifs may represent uncharacterized TFBS or serve as flanking sequences that fine-tune expression and facilitate TF binding^26,27^.

### Combinations of Novel and TFBS motifs are Co-Enriched in Regulatory Regions of the Top-Ranked T-cell-specific Genes

It is well established that cell-type-specific gene expression is primarily dictated by combinations of TFs that bind to DNA sequence motifs and work together to regulate gene expression^1–4^. Therefore, we asked whether there are specific pairs of motifs that are enriched in regulatory regions for the 22 top-ranked genes. To achieve this, we ran our statistical framework, this time using pairs of motifs consisting of a combination of two distinct JASPAR-annotated TFBS (n=900,601, TFBS:TFBS), or a combination of a known TFBS plus a novel motif (n=6,483,568, TFBS:Novel). In total, we identified 3,980 enriched TFBS:TFBS pairs that were enriched in regulatory regions for the 22 top-ranked T-cell genes and only 475 pairs for the control gene set containing the 22 bottom-ranked genes (**Figure 3A**). Of the pairs enriched in regulatory regions of the top-ranked genes, 90% (n=3,566) were enriched in distal regulatory regions, consistent with the notion that cell-type-specific gene expression profiles are primarily driven by combinations of sequence motifs within distal, rather than proximal regulatory regions. Further, 33% of the enriched pairs (n=1,352) are comprised of motifs where neither motif is enriched when considered alone (**Figure 3B**). These results are not unexpected given they are consistent with the notion that regulatory regions containing clusters of binding sites for TFs cooperate to control gene expression. As a specific example, 13 of the 22 top-ranked genes contained binding sites, within their distal regulatory regions, for FLI1 and GATA3, which are known to co-localize in T-cell regulatory regions^28^. Further, we observed 763,464 novel motif-containing pairs in the distal regulatory regions of the top-ranked genes, and only 14,682 such pairs in the distal regulatory regions of the bottom-ranked genes; a 52-fold difference (**Figure 3C**). Similarly, the number of enriched pairs containing a TFBS and a novel motif is larger in distal regulatory regions than in proximal regions. Overall, these findings suggest the possibility that known TFBS and novel motifs interact within distal regulatory regions to influence T-cell-specific gene expression.

**Figure 3.**
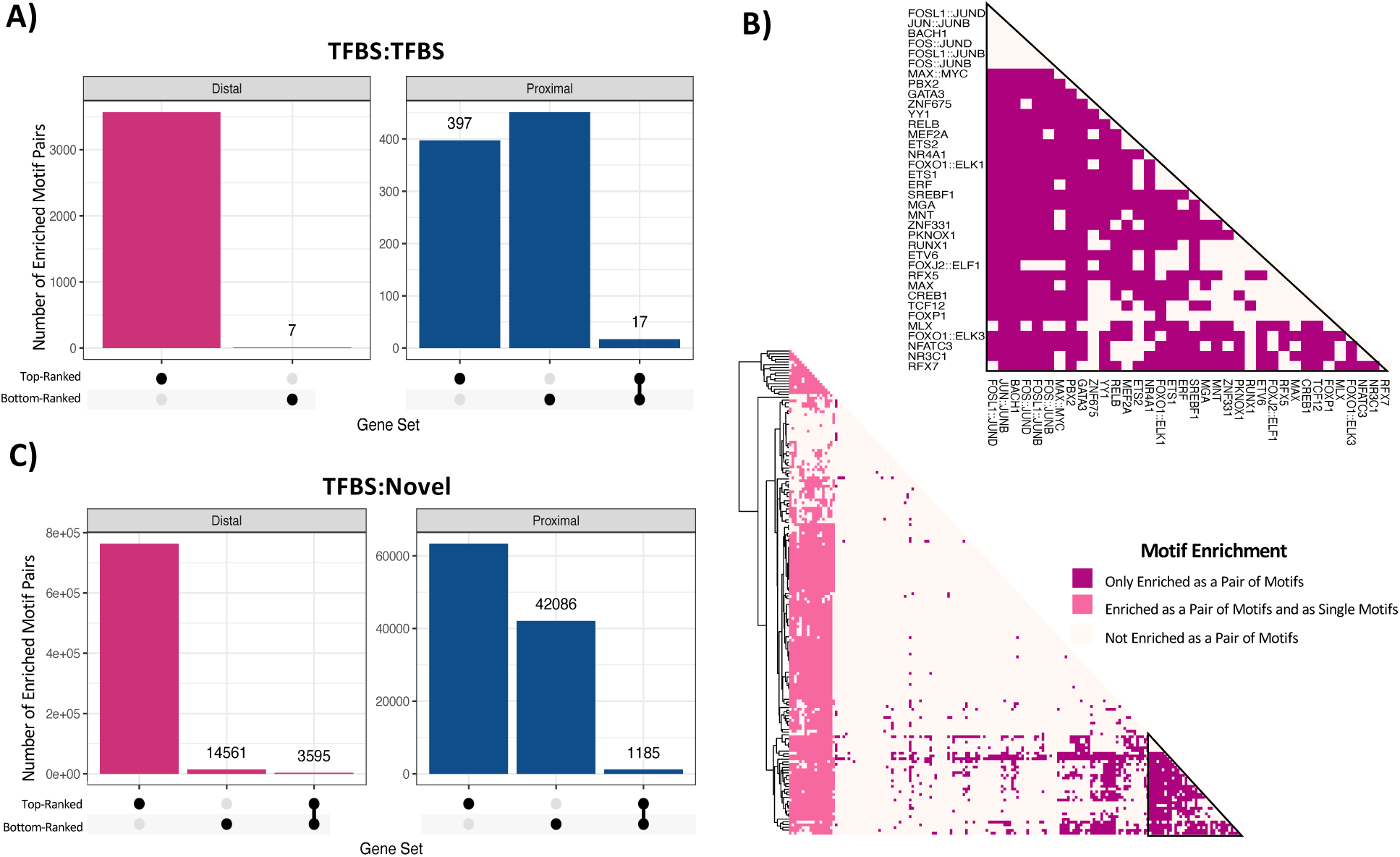
Representation of TFBS and novel motif combinations. A) The number of enriched motif pairs comprised of known TFBS from JASPAR (y-axis) by regulatory region type and the gene set (x-axis). B) Pairs of TFBS motifs in distal regulatory regions and their level of enrichment. Motif pairs are clustered based on their enrichment level. Motif pairs coloured pink are significantly enriched in regulatory regions for the top-ranked genes. Motif pairs in dark pink comprise two distinct TFBS motifs that are only enriched when considered as a pair of motifs. Motif pairs in light pink are enriched as a pair and contain one to two TFBS motifs that are also enriched when examined in isolation. C) The number of enriched motif pairs comprised of a TFBS and a novel motif (y-axis) by regulatory region type and gene set (x-axis).

### STARR-seq Library

To assess motif functionality, we selected 18 motifs (**Figure 4A, Supplemental Figure S6**) to evaluate, comprehensively, by STARR-seq^8^, a high-throughput sequencing assay that measures the ability of a regulatory element to enhance its own transcription. The candidate motifs included nine binding sites for transcription factors with established roles in T-cell function, such as TCF7, FOXO1::ELK3, and BATF::JUN, alongside nine novel motifs. The motifs were selected for their degree of enrichment in the regulatory regions of the 22 top-ranked genes (see Methods). Previous studies have demonstrated that order, orientation, and copy number can significantly influence the effect motifs have on gene transcription^7,9,10^, thus we designed a comprehensive synthetic oligonucleotide library consisting of i) oligos with one, two, or three copies of a given motif, ii) oligos that contain all possible permutations of two motifs, and iii) oligos that contain all possible permutations of three distinct motifs (**Figure 4A**). Each motif was tested in the forward (template) and reverse (non-template) orientations for all categories. The number of motif candidates we could comprehensively assess was limited to 18 by the feasibility of large-scale array-based oligonucleotide synthesis.

**Figure 4.**
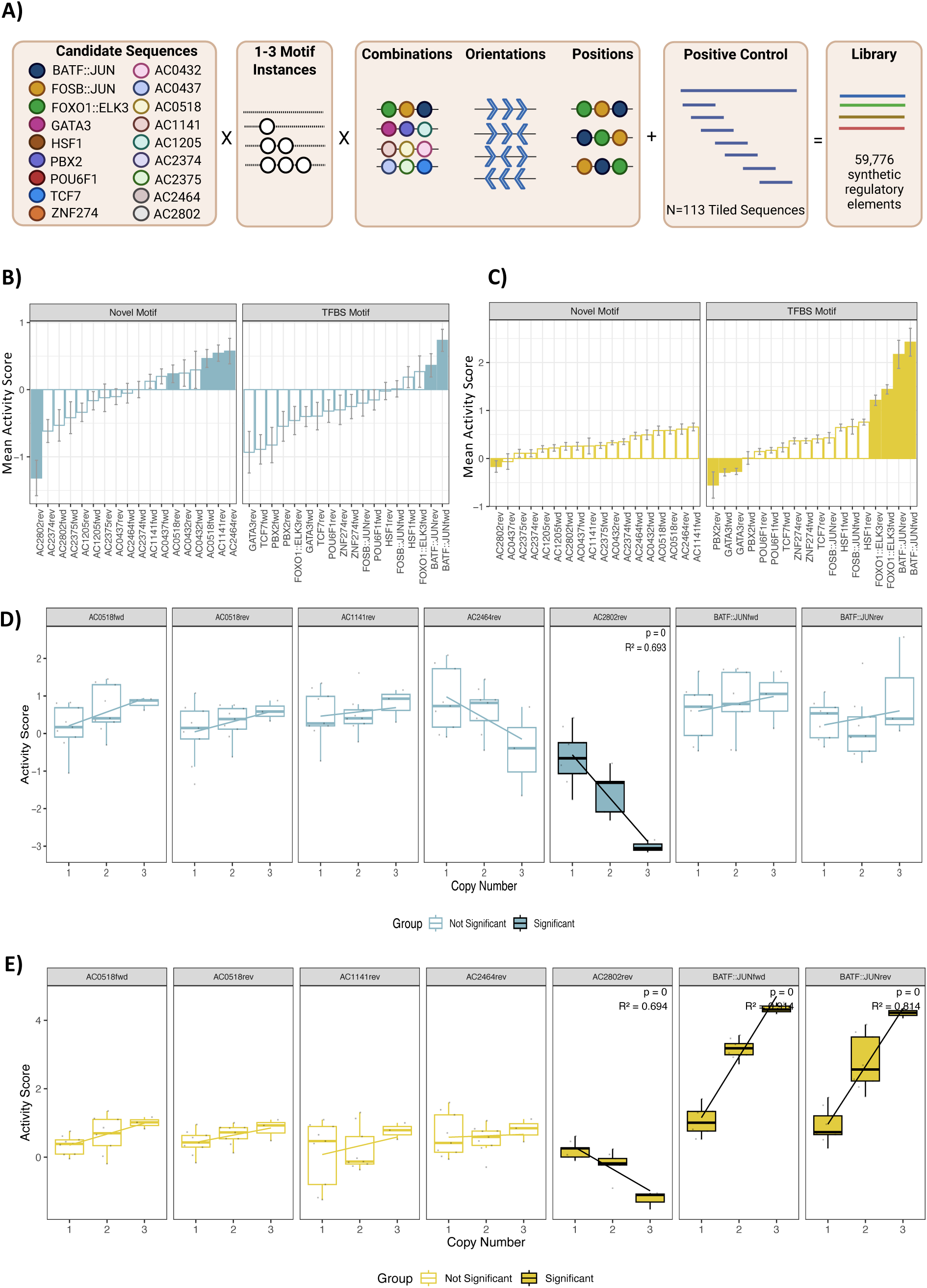
Novel motifs affect gene transcription in Jurkats. A) Shown, is a diagram of the STARR-seq oligo library design. The figure was adapted from Georgakopoulos-Soares et al^9^ under the Creative Commons license http://creativecommons.org/licenses/by/4.0/. Changes were made to the figure to account for the differences in our overall library design. Eighteen candidate motifs (nine novel and nine TFBS motifs) were embedded in a synthetic background DNA sequence. Each sequence contained one to three instances of a motif, separated by 10bp. The oligonucleotide library contains every possible combination, position, and orientation of the 18 candidate motifs. B) The mean activity score for the 18 motifs in their respective orientations for Jurkat and C) K562 cells are shown above. Mean activity scores were obtained from oligos containing the motif of interest only in its indicated orientation. Motifs with a significant effect on gene transcription were identified using a two-sided t-test that compared oligos containing the motif of interest to those that do not contain the motif. Only oligos containing 1-3 instances of the given motif in its respective orientation were used. Motifs with a significant effect on gene transcription are indicated as solid bars (Bonferroni corrected, p-value < 0.001, | Cohen’s D| > 0.5). Error bars represent the standard error of activity scores. D) Linear modelling was used to determine if there was an association between the number of copies of a motif and the mean activity score for the motifs with a significant effect on gene transcription in Jurkat cells and E) K562 cells (Bonferroni corrected, p-value < 0.007). Only oligos containing one to three copies of the motif of interest were included in the analysis.

To evaluate transcriptional activity, the oligonucleotide library was cloned into the STARR-seq reporter plasmid downstream of the core-promoter^29^, amplified, and transfected into two different host cell lines, Jurkat and K562, representing cells of T-cell and non-T-cell origin, respectively. Note that in order to evaluate candidate motifs in this manner, we needed to first embed them into a DNA background sequence. To reduce the chance of using a background DNA sequence with cryptic regulatory activity, we designed a synthetic background DNA sequence *in silico* using an iterative approach. This approach involved generating a random DNA sequence, screening the sequence for known TFBS and novel motifs, mutating any identified sites, and repeating the process until no motifs were found (**Supplemental Table S2**). Using flow cytometry, we confirmed that the synthetic background DNA sequence was transcriptionally quiet (**Supplemental Figure S7A**). Regarding the selection of a positive control DNA sequence, to our knowledge, no studies to date have reported a control DNA sequence with confirmed STARR-seq activity in Jurkat cells. Therefore, we selected a DNA sequence (seq1305, **Supplemental Table S2**) from a Lenti-MPRA-seq study that displayed strong enhancer activity in K562 cells as our positive control^30^. We verified the regulatory activity of this sequence using flow cytometry and observed strong GFP signals in both K562 and Jurkat cells (**Supplemental Figure S7B-C**). In total, we designed an oligonucleotide library comprised of 59,776 oligonucleotides comprising sets of motifs embedded within the validated synthetic background sequence. The positive control sequence (seq1305), which was initially 200bp, was tiled (n=113) into 88bp segments to match the same length as the synthesized oligos.

Initially, to determine the baseline representation of each oligonucleotide in our STARR-seq library, we sequenced the library directly. Comparing the normalised DNA counts (CPM) across the eight DNA replicates, we observed an average Pearson correlation of 0.957 (range 0.95-0.96, SD 0.0047, **Supplemental Figure S8**), indicating low technical variation. Additionally, we captured 99.9% (n=50,733) of our oligo library, demonstrating minimal loss of oligonucleotide representation during the process of cloning the oligos into the STARR-seq vector and amplifying the library. The library was then transfected into K562 and Jurkat cells in triplicate, using electroporation. RNA was isolated from the cells after 24 hours, and deep amplicon sequencing was performed using the Illumina platform. The correlation between the normalised RNA counts (CPM) across RNA replicates was lower in Jurkat cells (average: 0.55, range: 0.54-0.56, SD: 0.01, **Supplemental Figure S9A**) compared to K562 (average: 0.98, SD: 0.00, **Supplemental Figure S9B)**. We confirmed that library representation in Jurkat cells was not limited by sequencing depth, as each oligo had multiple reads (**Supplemental Figure S10**). Overall, we calculated activity scores (log2(RNA/DNA)) representing the level of enrichment between normalised RNA and DNA counts, for 50,634 and 50,650 oligos in Jurkat and K562 cells, respectively. Each oligo was required to have detectable RNA for at least two of the three RNA replicates. To account for the nominal effects of the background sequence, we subtracted the activity score for the “empty” background sequence containing no motifs from each oligo. The oligos derived from the positive control sequence led to increased gene transcription in K562 and Jurkat cells via STARR-seq (**Supplemental Figure S11**), consistent with the results we obtained by flow cytometry. The functional regions of the positive control sequences were towards the beginning and end of the full sequence.

### Novel Motifs Influence Gene Transcription in T cells

We asked whether there are novel motifs that can influence gene transcription preferentially in Jurkat cells. To answer this, we compared the activity scores in Jurkat cells between all oligos that contained the motif of interest to those that do not contain the motif. For this initial analysis, we only assessed oligos containing 1-3 instances of the given motif, and all motifs had to be in the same orientation. Overall, in Jurkat cells, five of the novel motifs influenced gene transcription, and in some cases, to an extent that was greater than the effect of known TFBS had on transcription (**Figure 4B**, two-sided t-test, Bonferroni corrected, p-value < 0.007, |Cohen’s D| > 0.5). Many effects were specific to the motif’s orientation, such as for AC2802, which only had a large repressive effect on transcription when in the reverse orientation. Four of the novel motifs had an activating effect on gene transcription, including AC0518 in any orientation, AC1141 in the forward orientation, and AC2464 in the reverse orientation. While AC2802rev had significant effects in Jurkat and K562 (**Figure 4B-C**), AC0518fwd/rev, AC1141fwd and AC2464rev only affected gene transcription in Jurkat cells, suggesting these motifs may affect gene transcription in a cell-type-specific manner. BATF::JUN, the only known TFBS with a significant effect on gene transcription in Jurkat cells, also exhibited activating effects in K562 cells (**Figure 1C**). The BATF::JUN TF is known to be functional in T cells and myeloid cells^31,32^.

### Copy Number and Position Influence the Motif’s Effects on Gene Transcription

For the motifs with a significant effect on gene transcription in Jurkats, we aimed to determine if there is an association between the number of motif copies and the level of gene transcription, a phenomenon that has been previously reported for TFBS^7,9,33^. Using linear modelling, we identified a novel motif, AC2802rev, where the variation in gene transcription could be partially explained by the number of motif copies (**Figure 4D**, Bonferroni corrected, p-value < 0.007). The negative association for AC2802rev was observed in both K562 and Jurkat cells, where nearly 70% of the variation in gene transcription levels could be explained by the number of copies of AC2802rev (**Figure 4D-E**). Finally, we explored whether the novel motifs have a different effect on gene transcription depending on whether it is the first or second motif in the oligo relative to the core promoter (**Supplemental Figure S12A)**. For this analysis, we obtained all oligos containing two distinct motifs and compared the activity scores for a motif of interest based on its position relative to the core promoter. We observed variable effects on gene transcription for three of the seven motifs in Jurkat cells (**Supplemental Figure S12B**, Wilcoxon t-test, Bonferroni corrected, p-value < 0.007). The largest positional preference was observed for AC2464rev, which either activates or represses gene transcription depending on its position. In addition to AC2464rev, AC0518rev also had significant positional preferences that were consistent in both cell lines (**Supplemental Figure S12C**), while AC2802rev had positional preferences that were specific to Jurkat cells. Overall, these results highlight that novel motifs can differentially affect gene transcription based on their position within a synthetic regulatory element in a cell-type-specific manner, irrespective of its neighbouring motifs.

### Pairs of Novel Motifs and known TFBS modulate Transcription

Next, we sought to determine whether specific pairs of motifs could synergistically activate or repress gene transcription more effectively than either of the motifs alone. First, we identified all pairs of motifs that have a significant effect on gene transcription. To achieve this, we compared the activity scores between oligos containing the two motifs of interest, to the oligos that do not contain the two motifs of interest. Only oligos containing 1-2 copies of two distinct motifs were used for this analysis. From this, we identified 62 and 199 motif pairs with effects on gene transcription in Jurkat and K562 cells, respectively (**Figure 5A**, two-sided t-test, Bonferroni corrected, p-value < 0.00008, |Cohen’s D| > 0.5). However, we hypothesized that many of these effects may be driven by an individual dominant motif rather than the pair of motifs working synergistically. Therefore, we required the Cohen’s D for the motif pair to be significantly greater than the Cohen’s D for the individual motif with the strongest effect on gene transcription (Z-score, p-value < 0.05). We also required the motif pair’s activity score to be greater than the sum of the individual motifs’ activity scores. Overall, we identified six pairs of motifs in Jurkats and one pair in K562 cells that exhibited activity levels greater than the activity level of either of the individual motifs in isolation (**Figure 5B, Supplemental Figure S13**) and hence not attributable to a single dominant motif. All of these motif pairs in Jurkat cells involved at least one novel motif, with AC2802 being the most prevalent. Notably, one pair was comprised of two novel motifs, AC0518rev and AC1141rev, and had the second-highest activating effect on gene transcription (**Supplemental Figure S13**). Although we observed cases where the addition of a second motif can alter gene transcription, these motifs appear to have an additive effect on gene transcription rather than a synergistic effect. Overall, these results highlight that gene transcription can be further modulated by the advantage of incorporating multiple distinct motifs in synthetic regulatory elements.

**Figure 5.**
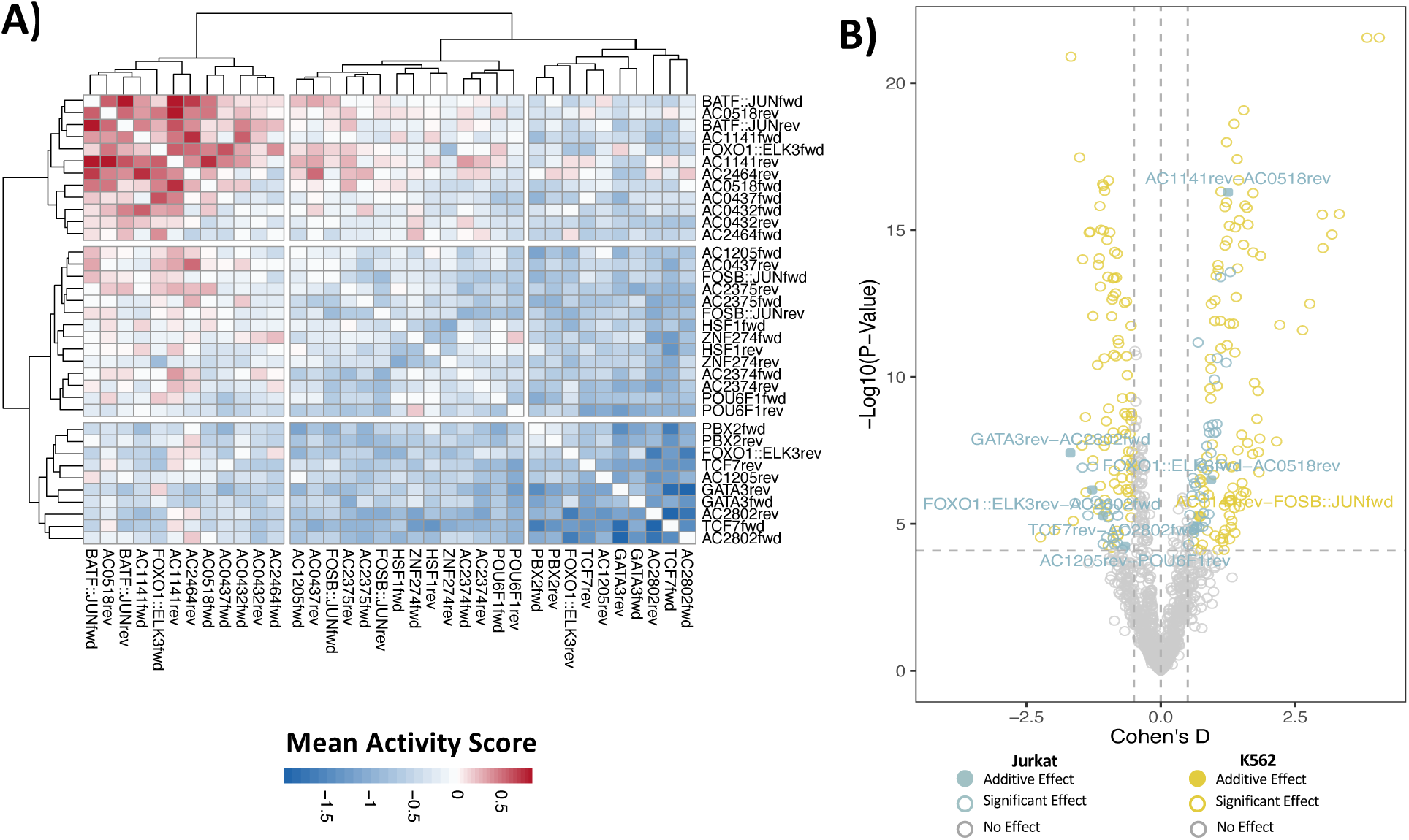
Pairs of motifs have effects on gene transcription in Jurkat cells. A) Heatmap showing the mean activity score for pairs of motifs for oligos containing one or two copies of two distinct motifs. B) Motif pairs with a significant effect on gene transcription (two-sided t-test, Bonferroni corrected, p-value < 0.00008, |Cohen’s D| > 0.5) are shown for Jurkat (blue) and K562 (yellow). Motif pairs that have a greater effect on gene transcription than either of the motifs in isolation are shown as solid points (mean activity motif pair > mean activity motif 1 + mean activity motif 2). Motifs within the pair are listed in alphabetical order. Motif pairs that do not have a significant effect on gene transcription are shown in grey. The dashed lines indicate the Cohen’s D threshold and p-value thresholds used to determine significance (|Cohen’s D| > 0.5, p-value > 0.00008).

### The Positional Preference of a Motif Depends on Its Neighbouring Motif

Finally, we explored whether the order of motifs within a specific pair can have a significant effect on gene transcription. For this analysis, we used oligos containing only one copy each of two distinct motifs and compared the activity scores for oligos containing the pair of motifs in one order compared to the opposite order (**Figure 6A**). In total, we identified 17 and 33 pairs of motifs for Jurkat and K562 cells where the effects on gene transcription differed according to the order the individual motifs within the oligo (**Figure 6B**, Wilcoxon t-test, Bonferroni corrected, p-value < 0.00008, |order1 activity score/order2 activity score|>2). For Jurkat cells, 82% (n=14) of the motif pairs involved a novel motif, eight of which were AC2464rev (**Figure 6C**). These results extend our findings described in the previous section, where we demonstrated that it is possible for a motif to have a preference for position, regardless of the secondary motif within the oligo (**Supplemental Figure S12C)**. Conversely, in K562 cells, only 63% (n=21) of the motif pairs involved a novel motif. Overall, these results highlight that the order of motifs within a synthetic regulatory element can have a significant effect on gene transcription, and that the effects can differ depending on the exact pair of motifs.

**Figure 6.**
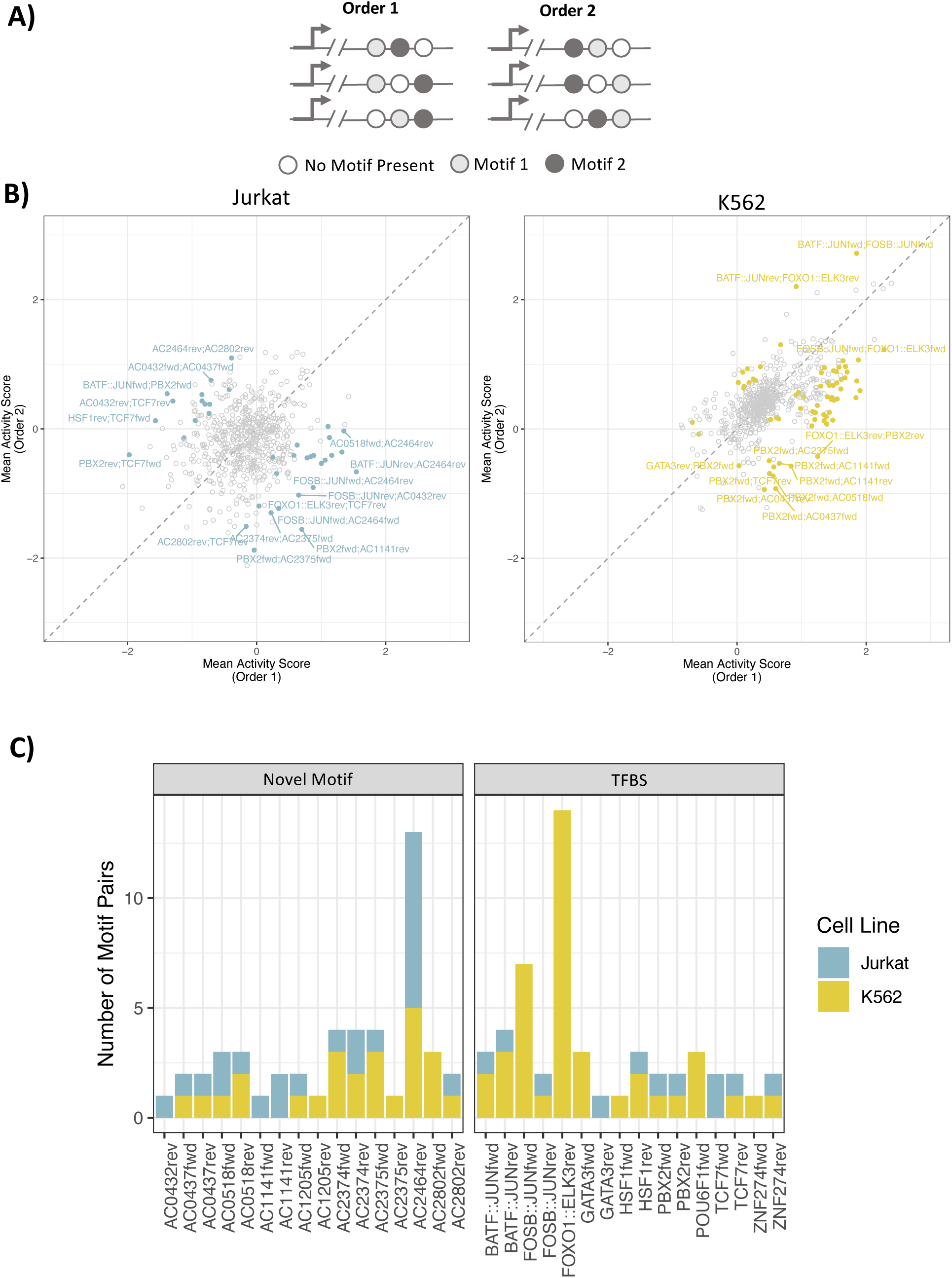
Motif pairs have varying effects on gene transcription depending on their order within synthetic regulatory elements. A) Schematic of the oligonucleotides used for the analysis. Only oligos containing a single copy of two distinct motifs were used. B) The mean activity score for oligos containing a single copy of each of the two distinct motifs is shown for Jurkat (left) and K562 cells (right). Pairs of motifs that have a different effect on gene transcription that is dependent on the order of the motifs are coloured blue (Jurkat) or yellow (K562). Grey points are pairs of motifs that did not meet the significance threshold. Tests were performed using the Wilcoxon t-test (Bonferroni corrected, p-value < 0.00008, |order1/order2|>2). C) The number of motif pairs containing a motif of interest that has a preference for order, as determined in Figure panel B.

## Discussion

We explored, by integrating existing multi-omics datasets, the sequence composition of regulatory regions of genes that are most selectively expressed in T cells. We demonstrated that these regions are enriched for binding sites for certain known transcription factors, but also, intriguingly, for DNA motifs we identified d*e novo*. Functional testing by STARR-seq revealed that half of the novel motifs modulated transcriptional activity at levels comparable to, or exceeding, those of known T-cell TFs, particularly in Jurkat cells. Many of these effects were orientation-dependent, further emphasizing the importance of motif directionality in transcriptional control. We also demonstrated that in some instances, increasing the motif copy number can amplify a motif’s effect on transcription, as demonstrated for the novel motif AC2802rev, which consistently repressed gene transcription in both Jurkat and K562 cells in a dose-dependent manner. Although we found that the majority of the candidate motifs operated independently, we were able to identify some specific pairs of motifs whose effects were additive. One of the strongest motif pairs comprised two novel motifs, AC1141rev and AC0518rev, and the influence of this pair on transcriptional activity was restricted to Jurkat cells. Lastly, we observed that novel motifs have strong preferences for a particular position within the synthetic regulatory element. AC2464rev, for example, had opposing effects on transcription that were dependent on its position in the oligo. In other cases, for example AC2464rev, the preferred position varied depending on what other motifs were present within the same element. Overall, these findings are consistent with the prevailing view of gene transcription, whereby combinations of motifs and their positioning form a regulatory “grammar” that governs cell-type-specific gene expression. Further, these results highlight the complexity of regulatory logic and underscore the advantages afforded by the use of comprehensive oligo libraries to systematically unravel this logic.

A number of previous studies have focused on identifying motifs enriched in T-cell regulatory regions^5,6,34–36^. Similar to our work, these previous studies were able to identify binding sites for TFs with known roles in T cells, including, for example, TFs from the FOS and JUN families^6,35,36^. Notably, some studies also reported enrichment of DNA binding sites for ubiquitous factors such as CTCF and SP1^35,36^ which suggests the methods used, while perhaps well suited to each study’s primary goals, may not have prioritized the identification of the motifs most relevant to T cells. Our work differs from these previous studies by using a gene-centric approach and by focusing on identifying motifs most strongly associated with T-cell-specific gene expression. Further, while two of the previous studies have reported novel T-cell-associated motifs^5,6^, these motifs were not assessed functionally. Finally, with one exception^34^, previous studies only focused on single motifs and did not search for combinations of enriched motifs, despite the knowledge that cell-type-specific gene regulation is primarily dictated by combinations of motifs and their associated factors. Thus, to our knowledge, ours is the only study to date to perform a comprehensive and unbiased search for enriched, previously uncharacterized DNA sequence motifs and exhaustively test resulting motif candidates for function in immortalized T cells.

In closing, we acknowledge that our work relied heavily on the work of other investigators who have generated extensive public data resources. Leveraging these resources, we identified new, previously undescribed sequence motifs that affect gene transcription in T cells, suggesting that currently annotated TFBS represent only a subset of motifs capable of modulating gene transcription in this important cell type. It is possible these novel motifs could provide binding sites for TFs (repressors, activators) or, in the case of motifs with repressive activity, induce structural changes in the DNA that impede the ability of the transcriptional machinery to bind the core promoter. The mechanism by which the novel motifs described herein may modulate transcription in the STARR-seq assay is unknown and will require further study. Nonetheless, our results highlight the need to consider novel motifs in gene regulation studies. Finally, our bioinformatic pipeline provides an approach for identifying motifs associated with cell-type-specific gene expression that could be applied to other cell types of interest, to probe other areas of biology.

## Methods

### Data Acquisition

Gene expression data from the Database of Immune Cell eQTLs, Expression, and Epigenomics^12^ used for the analyses presented in the current publication were downloaded from dbGaP phs001703.v1.p1 (Project #21785). Raw RNA-seq reads from normal and untreated samples were obtained from publicly available datasets (**Supplemental Table S1)** on the Sequence Read Archive (SRA). Annotated open chromatin regions in the hg19 genome were obtained from a publicly available dataset^13^ (https://zenodo.org/record/3838751#.ZAFOyezMJFw). Positional probability matrices (PPMs) for 949 human TFBS were downloaded from JASPAR^20^ (JASPAR2022_combined_matrices_2090_human_meme) using the search terms: species=*homo sapiens* and collection=*core*).

### Processing RNA-seq Data

Raw RNA-seq reads were trimmed to 50bp and adapters were removed using cutadapt (version 4.9, --length 50, --minimum-length 50, -a AGATCGGAAGAGCACACGTCTGAACTCCAGTCA, http://journal.embnet.org/index.php/embnetjournal/article/view/200/479). Processed reads were run through the GTEX V7 pipeline (https://github.com/broadinstitute/gtex-pipeline/tree/master/rnaseq) to generate gene expression values. Briefly, single-end reads were aligned to the human genome (hg19) using STAR^37^ (version 2.4.2a, -- overhang=49) and GENCODE v19 annotations. Only the forward reads were used for studies with paired-end reads. Transcripts were quantified with RSEM^38^ (version 1.2.22). RSEM estimated counts were normalized according to their library size using DEseq2^39^ (version 1.26.0) and adjusted for their effective gene length to enable comparisons across studies and genes. Final gene expression values were summarized at the cell-type level by calculating the median expression of the gene across samples for the same cell type.

### Processing H3K27Ac Hi-ChIP Interaction Data

Pre-processed H3K27ac Hi-ChIP data for CD4^+^ and CD8^+^ T cells were kindly provided by Dr. Ferhat Ay, Dr. Pandurangan Vijayanand, and Dr. Sourya Bhattacharyya^19^. Hi-ChIP interactions were generated following the methods described in Chandra et al^19^ using a bin size of 2.5kb. The CombineNearbyInteraction.py script from the FitHiChIP pipeline (https://github.com/ay-lab/FitHiChIP/tree/master/src) was used to merge adjacent significant Hi-ChIP interactions with scores exceeding 0.5. In summary, the curated dataset includes interactions for T-cells, where each interaction consists of two distinct genomic bins.

### Annotating Proximal and Distal Regulatory Regions

Using an existing set of annotated regulatory regions in T cells^13^, we curated a set of robust T-cell regulatory regions. Briefly, each regulatory region had to be annotated in at least one CD4^+^ T cell (Th1, Th17, CD4+, Th2, regulatory CD4^+^ T cells) and one CD8^+^ T-cell sample. Regulatory regions that overlapped by at least one base pair were merged into a single regulatory region. Regulatory regions were classified as proximal if they were within 500bp of a gene’s (protein-coding, TR_C_gene, IG_C_gene) transcription start site (TSS). All remaining regulatory regions were designated as distal. Regulatory regions on the Y or mitochondrial chromosomes were excluded from the analysis.

### Linking Distal Regulatory Regions to Putative Target Genes

We used the annotated interaction data and the curated T-cell regulatory region annotations to link distal regulatory regions to their target gene(s). First, for each gene, we obtained the coordinates of its proximal regulatory region(s). Next, we obtained all interactions with a bin overlapping the gene’s proximal regulatory region(s). Distal regulatory regions within the same or opposing bins for a given interaction were assigned as distal regulatory regions for the respective gene. For genes lacking merged interactions, we retained the original unmerged annotated interactions.

### Gene Lists

For each protein-coding, TR_C_gene, and IG_C_gene (excluding those on the Y and mitochondrial chromosomes), we calculated its aggregate expression value in T cells and non-T cells, separately by summing the expression values across all T cell and non-T cell samples. To ensure we were focusing on genes exhibiting adequate expression in T cells, we removed any genes whose aggregate expression value was below the median T-cell aggregate value (237.04). Using these aggregate expression values, we ranked each gene from high to low in T cells and low to high in non-T cells. Finally, a specificity score was calculated by ranking the sum of the two ranks, where small specificity score values represent genes exhibiting T-cell-specific expression. Lastly, we established a classification threshold based on the distribution of the specificity scores (μ = 10,033.5, σ = 2,134.1). Genes with a specificity score less than the mean specificity score minus four standard deviations were classified as the top-ranked T-cell genes (n = 22), while those with a score higher than the mean specificity score minus three standard deviations were designated as a comparator set of T-cell genes (n = 8,562). A control gene set of the bottom-ranked genes (n = 22) was also created.

### Novel Motifs

The genomic sequences of the regulatory regions linked to the 22 top-ranked T-cell genes were obtained using bedtools (version 2.30). Each regulatory region was only recorded once to avoid double-counting. The sequences were broken down into k-mers (k=12) and clustered into groups based on a Hamming Distance of one. Each group represents a set of k-mers where each k-mer differs by at most one nucleotide from at least one other k-mer in the group. PPMs were generated for each group, and TOMTOM^40^ (version 5.5.2) was used with default settings to remove PPMs with a match to TFBS motifs from the JASPAR database^20^ (JASPAR2022_combined_matrices_2090_human_meme).

### Identifying Enriched Motifs

We developed a statistical framework in Python to assess the enrichment of motifs in regulatory regions associated with genes of interest. Briefly, FIMO^41^ (version 5.4.1, --thresh 0.0001) was used to find occurrences of novel motifs and TFBS (motifs from JASPAR^20^). Only matches with a score greater than 11.16 were used for analysis (the mean score for all predicted motifs in proximal and distal regulatory regions associated with the 22 top-ranked T-cell genes). For each motif, we quantified the number of the top-ranked T-cell genes linked to at least one regulatory region containing the motif. To evaluate the statistical significance, we generated 10,000 size-matched sets of randomly sampled genes from the comparator set of T-cell genes. For each iteration, we calculated the number of genes linked to regulatory regions that contained the motif. The p-value was defined as the proportion of iterations where the number of genes from the top-ranked gene set exceeds the gene count for the number of randomly-sampled genes from the comparator gene set. In addition to individual motifs, we applied the same methodology to pairs of motifs, which could reside in the same or different regulatory regions associated with the same gene. For analyses, we only used TFBS for TFs that were expressed in T cells. TFBS for TFs with a T-cell aggregate expression value less than 237.04 were removed. Motifs were designated enriched if they had a p-value < 0.05 and a fold change > 2.

### Selecting Candidate Motifs and Representative Sequences

To select candidate motifs for experimental validation using the STARR-seq assay, we focused on results for pairs of motifs containing two distinct TFBS (TFBS:TFBS) or a TFBS and a novel motif (TFBS:Novel). These two groups of motif pairs were ranked independently based on two metrics: 1) the number of genes from the top-ranked gene set and 2) the average number of genes from the comparator gene set linked to regulatory regions containing the motif pair. We then summed the two ranks to assign a final score for each pair. To select TFBS motifs, we stepped through the ranked TFBS:TFBS pair list. We extracted pairs until we obtained a collection of nine unique TFBS motifs, ensuring that no two motifs belonged to the same motif archetype as defined by Vierstra et al^3^ (https://resources.altius.org/~jvierstra/projects/motif-clustering-v2.0beta/). This step ensures each selected motif represents a distinct DNA-binding archetype. Next, to capture potential synergistic interactions between TFBS and novel motifs, we filtered the TFBS:Novel motif pairs to retain only those containing one of the previously selected TFBS motifs. From this filtered set, we selected the top nine ranked novel motifs for validation using the same approach described above. For the 18 selected motifs, we chose a single DNA sequence to represent each motif in the STARR-seq library. Using the predicted motif occurrences within regulatory regions for the top-ranked T-cell genes, as determined by FIMO, we selected the most prevalent natural sequence as our representative sequence. If there was a tie, we selected the sequence with the highest FIMO score.

### Background DNA Sequence

To test our candidate sequences using the STARR-seq assay, we needed to embed our candidates into a background sequence with minimal regulatory activity. We designed a synthetic background sequence *in silico* by generating a random sequence of 125 bp. We then used FIMO to check for the presence of any motifs (TFBS, uncharacterized motifs, or motifs associated with histone marks^42^, and selectively mutated any subsequences containing motifs. This process was repeated until we obtained a synthetic DNA sequence with no motifs. The sequences were then flanked with AgeI and SalI restriction sites to facilitate cloning into the STARR-seq vector and ordered as a gene segment using TWIST Biosciences (San Francisco, USA).

### Library Design

*In silico*, we designed oligonucleotide sequences containing up to three of our 18 candidate sequences embedded into a synthetic background sequence. Candidate sequences were either in the template or non-template orientation and were spaced 10bp apart. After designing the library, the oligonucleotide sequences were flanked by AgeI and SalI restriction sites to facilitate cloning into the STARR-seq vector (hSTARR-seq_ORI, Addgene #99296) and the necessary priming and Illumina adapter sequences were added to enable sequencing (**Supplemental Table S3**). Lastly, as a positive control sequence, we used seq1305(+) (chr1:61048980-61049180); a 200 bp DNA sequence previously determined to have strong enhancer activity in K562 cells^30^. To match the length of our synthesized oligos, this sequence was tiled into every possible 88bp segment, resulting in 113 unique sequences. In total, the final oligonucleotide library comprised 59,776 distinct sequences and was synthesized by TWIST Biosciences (San Francisco, USA).

### Plasmid Production, Transfection, and Flow Cytometry

For testing the baseline regulatory activity of our synthetic background DNA sequence, the non-functional truncated GFP (trGFP) in the hSTARR-seq_ORI vector from Addgene (plasmid # 99296) was replaced with a full-length GFP sequence using AflI and AgeI (hSTARR-seq_ORI_fullGFP). The synthetic background DNA sequence was cloned into hSTARR-seq_ORI_fullGFP using AgeI and SalI. A transfection control plasmid containing mStrawberry was used to assess transfection efficiencies. Jurkat and K562 cells were co-transfected with the two plasmids using the Neon transfection system using 100uL tips with Jurkat using 5ug of DNA at 2x10^7^ cells/mL, pulse voltage 1350v, 10ms pulse width and 3 pulses, and K562 using 5ug of DNA at 1.5x10^7^ cells/mL, pulse voltage 1350v, 10ms pulse width and 4 pulses. Cells were assessed for GFP positivity via flow cytometry.

### Vector Preparation

hSTARR-seq_ORI vector from Addgene (plasmid#99296) was digested using AgeI-HF and SalII-HF using the following conditions: 5ug DNA with 500 units of AgeI-HF (NEB) and 500 units of SalI-HF (NEB) in a 500uL reaction incubated overnight and then heat-inactivated at 65°C for 20 minutes. Digested DNA was gel-purified to separate digested linear vectors and fragments recovered using the Monarch Spin DNA Gel Extraction kit as per the manufacturer’s instructions.

### Library Amplification

Four PCR amplification reactions were performed using the KAPA HiFi DNA Polymerase kit with the conditions mentioned below. PCR reactions were cleaned up using the QIAquick PCR purification kit following the manufacturer’s instructions with the following modifications: DNA was eluted in 50µl EB, re-applied to the column and eluted again. 50ng adapter-ligated ssDNA oligo, resuspended at 10ng/ul was used with 2.5uL of both the SS_oligo_f (10uM), and SS_oligo_r (10uM) in a 50uL KAPA HiFi PCR reaction. PCR reaction was 1 cycle: 98°C for 45s, 10 cycles: 98°C for 15s, 60°C for 30s, 72°C for 45s, 1 cycle: 72°C for 120s. Primer sequences are available in **Supplemental Table S3**.

### Library Cloning

Purified and PCR-amplified library inserts were cloned into the hSTARR ORI-seq plasmid using the conditions listed below. A ligation reaction using 125ng of the AgeI/SalI digested hSTARR-seq ORI vector with 16.67ng of the purified library insert in a 10μL reaction with T4 DNA ligase (NEB). Ligation was run at 16°C in a thermal cycler for 18 hours. A total of four cloning reactions were performed. All four cloning reactions were pooled up to 100μL and purified using the Qiagen MinElute column according to the manufacturer’s instructions and eluted into 12.5μL of EB.

### Transformation

Electrocompetent MegaX DH10B bacteria were transformed into four transformations for our library. 2.5μL of the purified and pooled library from the library cloning is added to pre-cooled 1.5mL Eppendorf tubes. 20μL of MegaX DH10B bacteria is added to each reaction tube and transferred to pre-cooled 1mm MicroPulser cuvettes (BioRad). Electroporation of each bacteria-DNA mix was done at 2kV, 25uF, 200*Ω,* and immediately added pre-warmed recovery medium to the cuvette and transferred to 14mL polypropylene round-bottom tubes for incubation for 1hr at 37°C with shaking. The four transformation reactions are pooled and plated on LB/Carbenicillin plates at 1:10, 1:50, 1:500, and 1:5000 dilutions to estimate library complexity and equal volumes of the pooled transformation reactions were also added to four 500mL LB/Carbenicillin Erlenmeyer flasks to amplify by overnight culture at 37°C. Bacteria cultures are harvested and resuspended bacteria are pooled, and distributed into four 50mL Falcon tubes and the bacterial pellet weight was determined. DNA from bacterial pellets were prepared using the PureLink HiPure Expi Plasmid Gigaprep kit (Thermofisher) as per the manufacturer’s protocol and eluted into ddH_2_O.

### UMI PCR and Sequencing Ready PCR, Azenta Sequencing

1ug of DNA of the plasmid library was digested with AgeI-HF (NEB) overnight and then purified by QIAquick PCR purification kit following the manufacturer’s instructions and eluted in 35μL EB. 18μL of the linearized library DNA was used for the UMI PCR: 7.5μL SS_UMI_r primer (10μM), 50μL KAPA 2x HIFI Hotstart ready mix in a total of 100μL reaction. One cycle of 98°C for 60s, 65°C for 30s, and 72°C for 90s to integrate UMI into the STARR-seq library and then purified using QIAQuick PCR purification kit as per the manufacturer’s instructions, except that the columns were eluted twice with 10μL EB.

Two sequencing-ready PCR reactions were carried out using 100ng of UMI-tailed template DNA, 2.5μL SS_i5_XX_f primer (10uM), 2.5μL SS_P7seq_r primer (10uM), 25μL KAPA 2x HiFi HotStart Ready Mix in 50μL total reaction. PCR conditions were: 1 cycle at 98°C for 45s, 9 cycles at 98°C for 15s, 60°C for 30s, 72°C for 45s, 1 cycle at 72°C for 120s. PCR reactions are purified using the QIAQuick PCR purification kit as per the manufacturer’s instructions, using only one column and eluting twice with 10μL EB and pooled. Primer sequences are available in **Supplemental Table S3**. 150bp paired-end reads were generated using an Illumina Novaseq machine.

### Cell Lines and Transfections

Jurkat (gifted from the Weng lab) and K562 (ATCC CCL-243) cells were cultured in cRPMI media (RPMI media with 10% FBS, 1% Penicillin/streptomycin, 1mM HEPES, 1mM sodium pyruvate) and verified by ATCC STR profiling and tested to be mycoplasma negative using the VenorGeM mycoplasma detection kit (Sigma-Aldrich). Cells were co-transfected using the Neon transfection system using 100μL tips with Jurkat using 5ug of DNA at 2x10^7^ cells/mL, pulse voltage 1350v, 10ms pulse width and 3 pulses, and K562 using 5ug of DNA at 1.5x10^7^ cells/mL, pulse voltage 1350v, 10ms pulse width and 4 pulses. Transfections were done in triplicate. Cells were harvested 24 hours after transfection, washed in PBS to remove growth media and cells pelleted. RNA was prepared using the Qiagen RNeasy Maxi Kit as per the manufacturer’s instructions.

### cDNA Synthesis, Purification, Library Construction, and Sequencing

We generally followed the protocol by Neumayr et al. (2019). The quality and quantity of total RNA were determined using RNA Nano kits (Agilent). mRNA/poly(A)+ RNA was extracted using the NEBNext High-Input Poly(A) mRNA Magnetic Isolation Module (NEB, USA) following the manufacturer’s instructions and treated with DNase 1 to eliminate DNA contamination. cDNA was generated from each sample using a Maxima H kit (ThermoFischer Scientific, see detailed method below). We then performed junction PCR (jPCR) as described in Neumayr et al. (2019). Briefly, a 50 µL PCR reaction was prepared containing 20 µL of cDNA, 25 µL of LongAmp 2x Master mix (NEB), and 2.5 µL of each of the SS_intspan_f and SS_P7seq_r 10 µM primers. PCR was performed with initial denaturation at 98 °C for 60 seconds (s), and 16 cycles of denaturation at 98 °C for 15 s, annealing at 65 °C for 30 s, and extension at 72 °C for 70 s, and a final extension at 72 °C for 300 s. The products of jPCR were purified and SPRI beads and quantified using Qubit 1x dsDNA HS reagent (Thermo Fischer Scientific). A final sequencing-ready PCR was performed to add Illumina sequencing adapters to the fragments. Briefly, a 50 µL PCR reaction was prepared containing 20 µL of jPCR products, 25 µL of LongAmp 2x Master mix, and 2.5 µL of each of SS_i5_f and SS_P7seq_r primers at 10 µM. PCR was performed with initial denaturation at 98°C for 45s, 5 cycles of denaturation at 98°C for 15s, annealing at 65°C for 30s, extension at 72°C for 45s, and a final extension at 72°C for 120s. The final sequencing PCR products were purified using SPRI beads and quantified using Qubit 1x dsDNA HS reagent. Samples were pooled and sequenced on an Illumina Novaseq X Plus machine. Primer sequences are available in **Supplemental Table S3.**

### Processing STARR-seq Sequencing Data

Oligos on average had an average depth of 432 read pairs across the eight libraries (**Supplemental Table S4**). For amplicon libraries, oligos had an average depth of 60 and 177 read pairs for Jurkat (n=3) and K552 cells (n=3), respectively (**Supplemental Table S4**). The 10bp UMIs were extracted from the sequencing reads using umi-tools^43^ (version 1.1.4, bc-pattern=.*AGATCGGAAGAGCACACGTCTGAACTCCAGTCAC(?P<UMI_1>.{{10}}).*). Illumina P7 and P5 adapters (**Supplemental Table S3**) were removed using cutadapt (version 4.7, quality_cutoff=10, min_length=80). Reads were mapped to our oligo library using bowtie2^44^ (version 2.5.3). Only the primary alignments were retained, as well as properly paired reads where both reads had a mismatch of 0 with the aligned sequence (samtools version 1.19 -F 780, -f 2, bamtools 2.5.2, NM:0). The number of unique UMIs observed for each oligo was calculated to obtain a DNA and RNA count.

### STARR-seq Analysis

Analysis of the STARR-seq data was performed for K562 and Jurkat cells separately. Only oligos with a DNA count >= 10 across all eight replicates were retained. Oligos with an RNA count of 0 in all replicates were also removed. RNA and DNA counts were adjusted for library size using counts per million (CPM). Finally, the DNA replicates were collapsed into a single value by averaging the DNA CPM values. For oligos with an RNA and DNA count greater than zero, an activity score representing the log2(RNA/DNA) was calculated for each replicate (n=3). The activity score for the synthetic background oligo without any motifs was subtracted from the activity score for each oligo to obtain the final, adjusted activity score.

## Data Availability

The STARR-seq data generated in this study have been deposited in GEO under GSE301996. Source data is provided with the paper and in the GitHub repository at https://github.com/nknoetze/tcellregulatorylogic.

## Code Availability

All code is publicly available at the following GitHub link: https://github.com/nknoetze/tcellregulatorylogic.

## Author Contributions

R.H. and N.K. conceived the study. N.K. and S.D.B. performed the analyses. N.K. and S.D.B. designed the oligo library. E.Y. performed the flow cytometry and STARR-seq experiments with help from Canada’s Michael Smith Genome Sciences Centre. N.K. wrote the manuscript with input from the authors. All authors reviewed and approved the final manuscript.

## Acknowledgements

We acknowledge the Genomics Platform at Canada’s Michael Smith Genome Sciences Centre (GSC), Provincial Health Services Authority, especially Yongjun Zhao and Simon Haile Merhu for their technical input and beneficial discussion, and Andrew Mungall and Richard Moore for the support in sample processing and sequencing, as well as the GSC’s Collaborative Services Team. This research was funded by the Canadian Institutes of Health Research (PJT-175287).

## Supplemental Figures and Tables

**Supplemental Figure S1.**
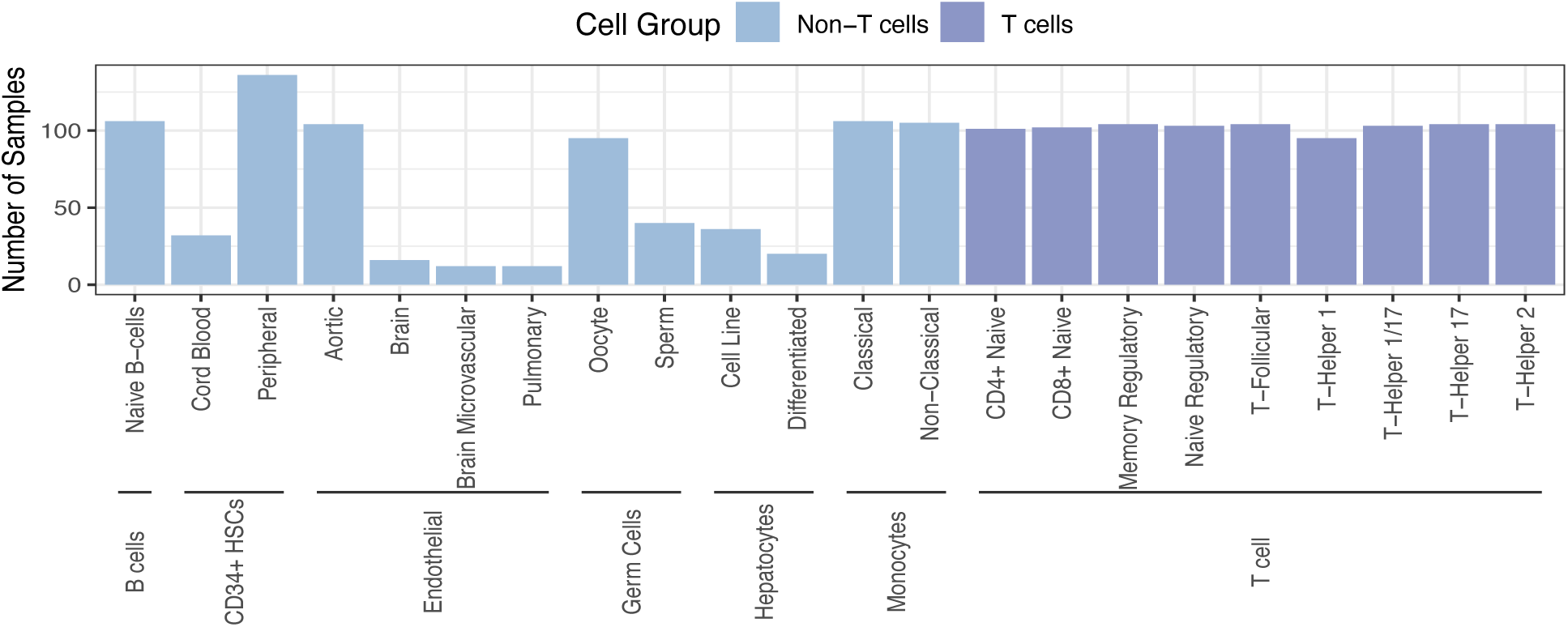
Number of RNA-seq samples used for gene expression analyses. RNA-seq data from the DICE consortium^12^ and the SRA (see Supplemental Table S1) were used to identify protein-coding genes expressed in T cells. The plot displays the number of RNA-seq samples used to calculate the median expression for each protein-coding gene in the genome across each cell type independently (y-axis).

**Supplemental Table S1.** Bioproject accessions for the RNA-seq datasets obtained from the SRA. PRJNA494278 was obtained from the dbGaP website, under phs001703.v1.p1 (Project #21785). The table includes the cell type(s) for each dataset and the number of samples used.

| Cell Group | Study | Cell Type | Number of Samples |
| --- | --- | --- | --- |
| T cell | PRJNA494278 | Naive CD4<sup>+</sup> T cells | 101 |
|  |  | Naive CD8<sup>+</sup> T cells | 102 |
|  |  | Follicular Helper T cells | 104 |
|  |  | Th1 | 95 |
|  |  | Th1-17 | 103 |
|  |  | Th17 | 104 |
|  |  | Th2 | 104 |
|  |  | Memory Regulatory T cells | 104 |
|  |  | Naive Regulatory T cells | 103 |
|  |  | Naive B cells | 106 |
|  |  | Classical Monocytes | 106 |
|  |  | Non-Classical Monocytes | 105 |
| Non-T cell | PRJNA687693 | Brain microvascular endothelial cells | 12 |
|  |  | Pulmonary endothelial cells | 12 |
|  | PRJNA515044 | Brain endothelial cells | 16 |
|  | PRJNA579487 | Aortic endothelial cells | 104 |
|  | PRJNA431399 | Terminally differentiated hepatic cells | 20 |
|  | PRJNA622787 | Hepatocyte cell line | 36 |
|  | PRJNA726552 | Sperm cells | 6 |
|  | PRJNA310976 |  | 34 |
|  | PRJNA666614 | Oocyte cells | 75 |
|  | PRJNA484890 |  | 20 |
|  | PRJNA211925 | CD34<sup>+</sup> CD45RA<sup>-</sup> hematopoietic stem cells from cord blood | 136 |
|  | PRJNA419347 | CD34<sup>+</sup> peripheral hematopoietic stem cells | 32 |

**Supplemental Figure S2.**
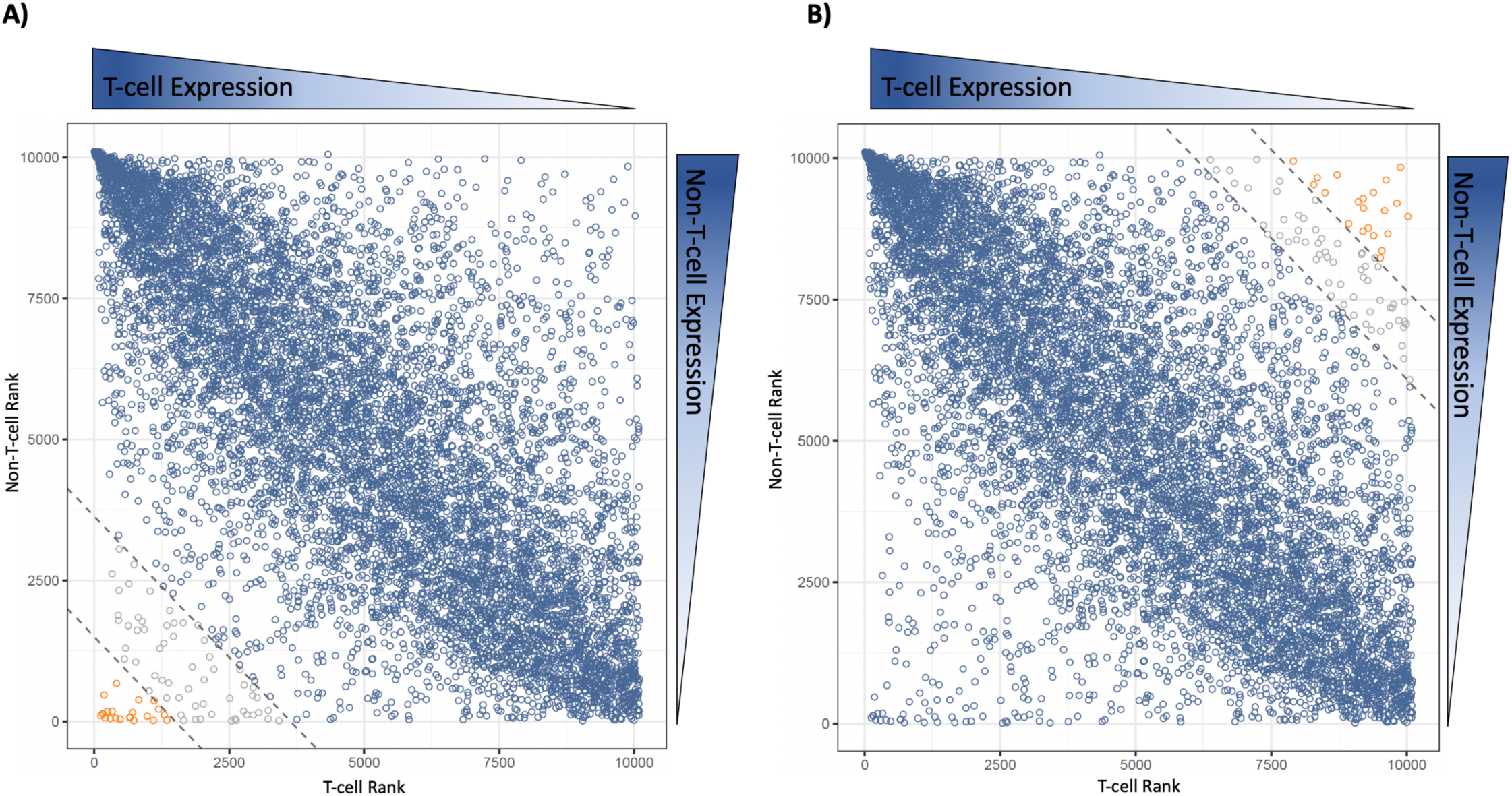
The relationship between gene ranks for T-cells and non-T-cells. The plots display the ranks for all genes expressed in T cells. Genes with higher expression in T cells receive a higher T-cell rank (x-axis), and genes with lower expression in non-T cells receive a higher non-T-cell rank (y-axis). A) The top-ranked T-cell genes (n=22, shown in orange) have a specificity score that is more than four standard deviations from the mean. The comparator gene set containing the non-specific T-cell genes (n=8,562, shown in blue) have specificity scores within three standard deviations from the mean. B) We created a size-matched control gene set by selecting the 22 bottom-ranked genes as our genes of interest (orange) and the top 8,562 ranked T-cell genes as our comparator set of genes (blue).

**Supplemental Figure S3.**
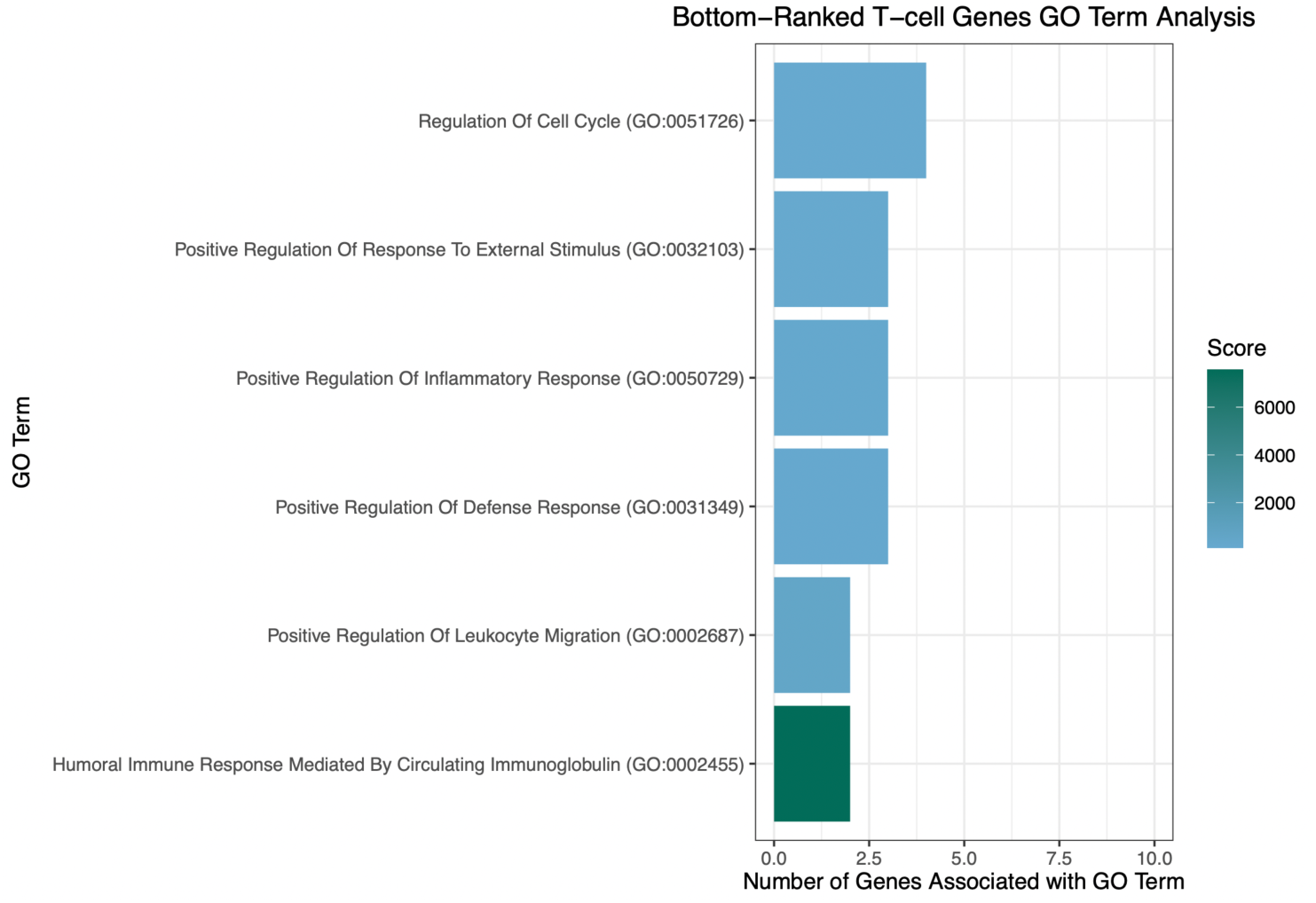
The 22 bottom-ranked T-cell genes are associated with immune-related biological processes. The bar plot shows the enriched GO terms (y-axis, Benjamini-Hochberg corrected, p-value < 0.05) for the 22 bottom-ranked T-cell genes. The bars correspond to the number of genes associated with the specified GO Term. Colour corresponds to the score obtained from the GO term analysis. GO terms are ordered by their adjusted p-value.

**Supplemental Figure S4.**
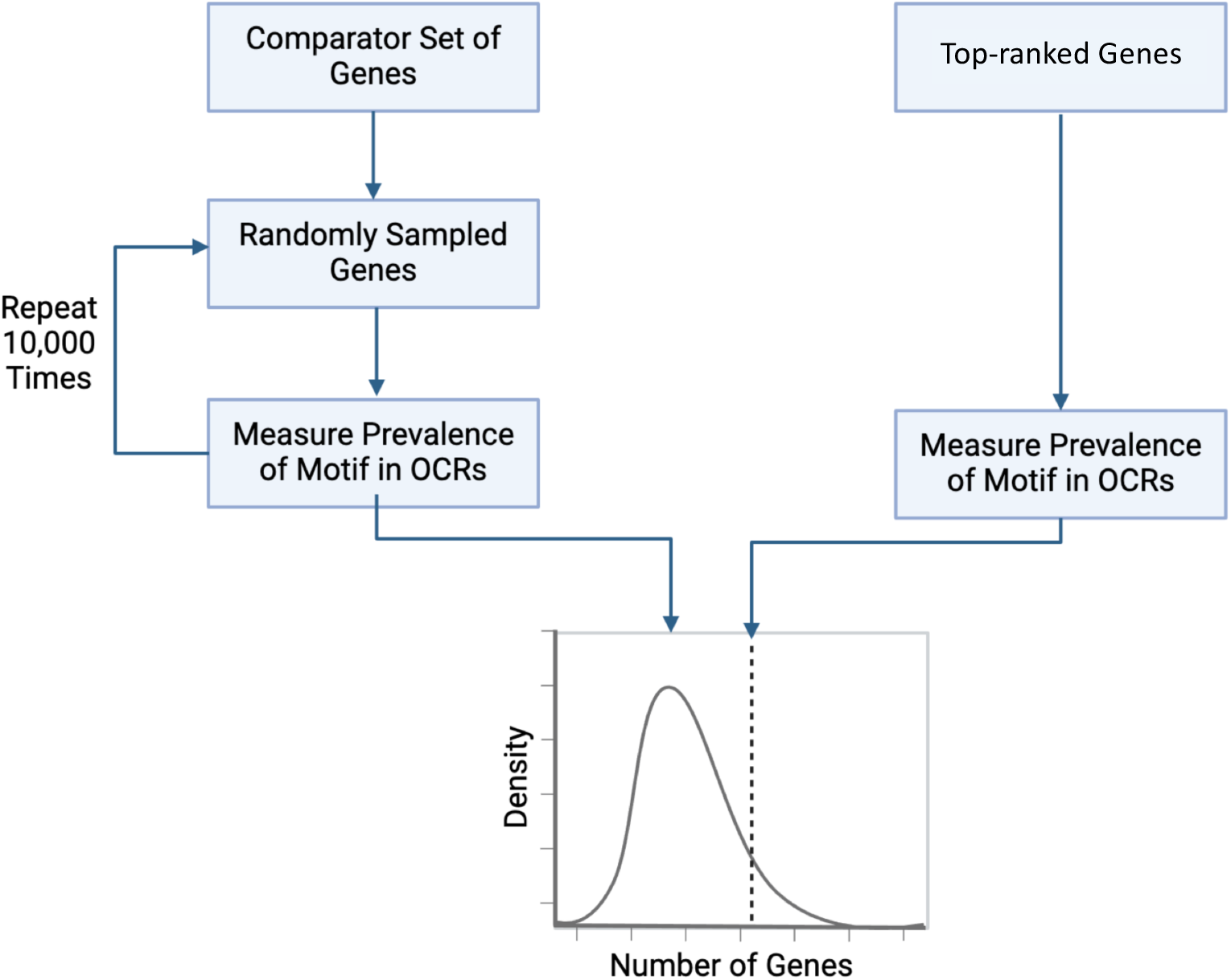
Illustration of the empirical framework used to identify enriched motifs. The framework utilizes a set of genes of interest (top-ranked T-cell genes) and a set of comparator genes (non-specific T-cell genes). For the genes of interest, the OCRs are obtained, and FIMO is used to identify sites within the OCRs corresponding to TFBS motifs or novel motifs. The framework then obtains 10,000 sets of genes of the same length as the genes of interest and repeats the process. Motifs are enriched if less than 5% of the random samplings exhibited a level of enrichment as large as the level observed for the genes of interest. Additionally, there needs to be at least twice as many genes of interest with the motif compared to the comparator set of genes.

**Supplemental Figure S5.**
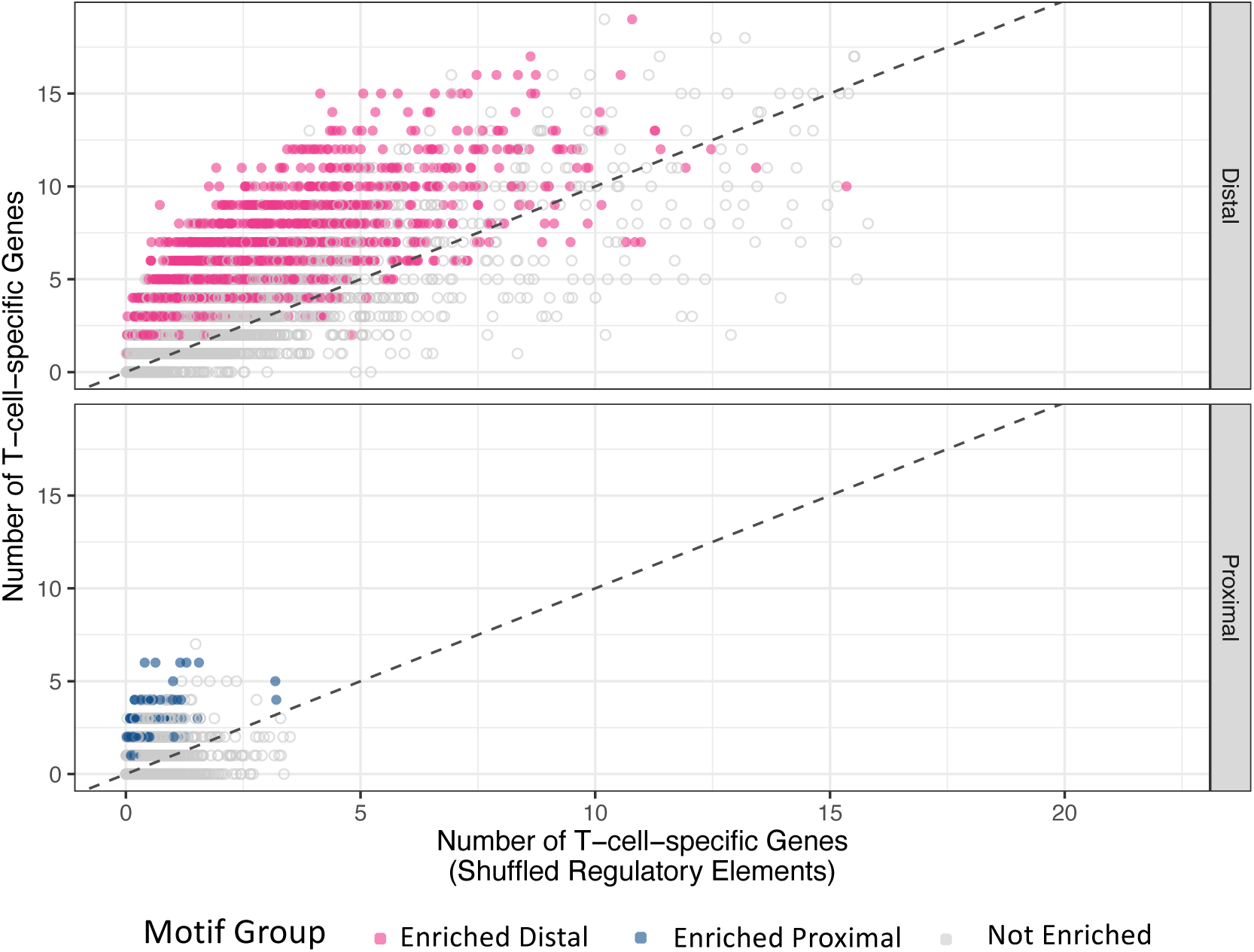
The statistical framework was used to identify enriched novel motifs in distal (pink) and proximal (blue) regulatory regions. Non-enriched motifs are shown in grey. FIMO was used to identify sites for all novel motifs in regulatory regions for the 22 top-ranked T-cell genes (y-axis). The regulatory regions for the 22 top-ranked T-cell genes were randomly repositioned across the genome 10,000 times to obtain the expected number of motif sites in the genome (x-axis).

**Supplemental Figure S6.**
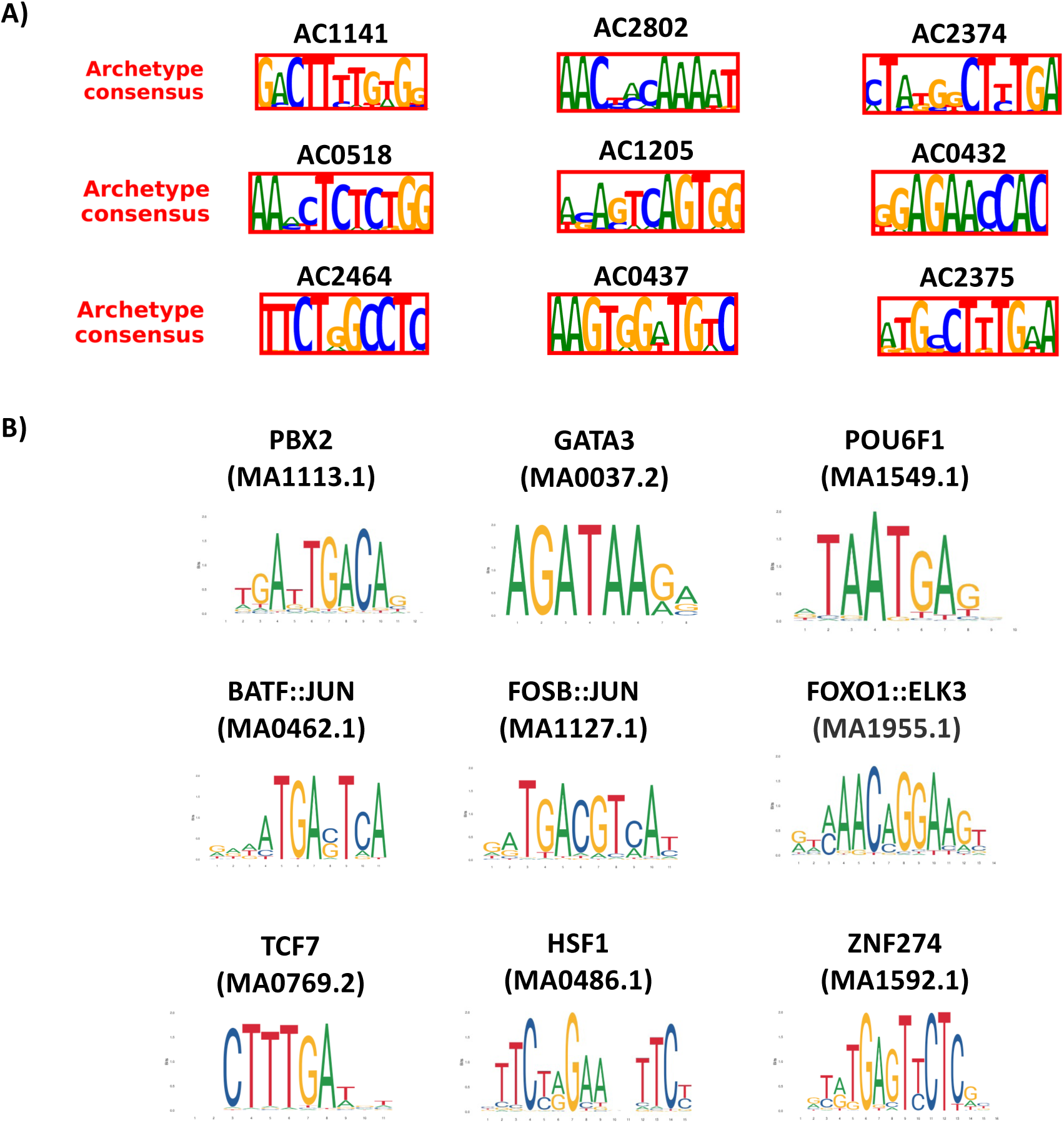
Candidate novel motifs and TFBS selected for validation using the STARR-seq assay. A) Novel motifs and B) TFBS were selected for their enrichment in regulatory regions for the top-ranked T-cell genes (see Methods).

**Supplemental Figure S7.**
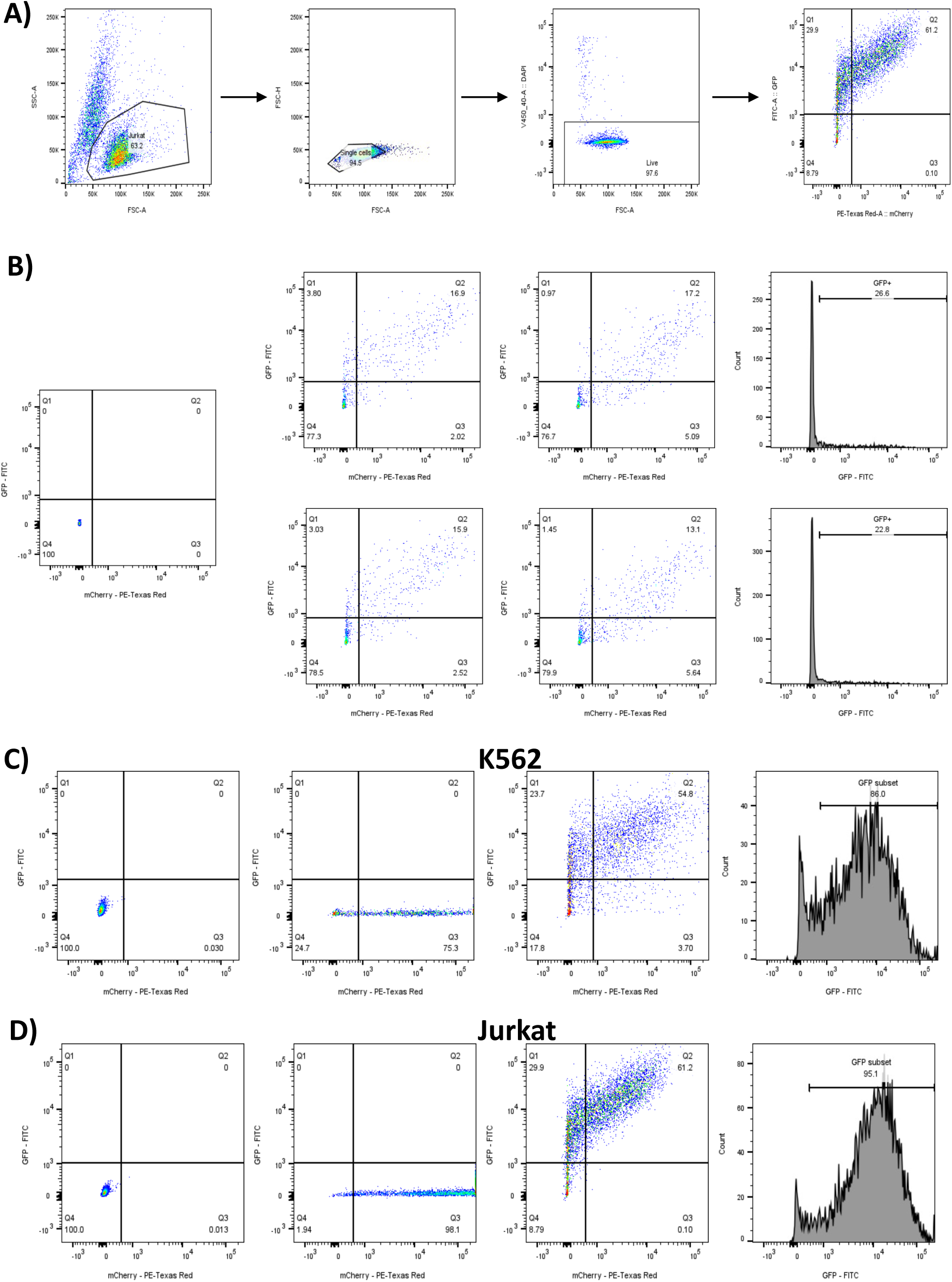
Flow Cytometry results for the positive control and synthetic background sequence. Jurkat and K562 cells were transfected with the STARR-seq vector containing the synthetic background DNA sequence, or the positive control sequence (seq1305), alongside a vector containing mCherry as a transfection control. Transfections were successful for both cell lines, and GFP expression was detected using flow cytometry. A) Gating strategy used in the flow cytometry experiment. B) The synthetic DNA sequence designed *in silico* had minimal expression in Jurkat cells. The positive control sequence drove GFP expression at higher levels than the synthetic background sequence in both C) K562 and D) Jurkat cells.

**Supplemental Table S2.**
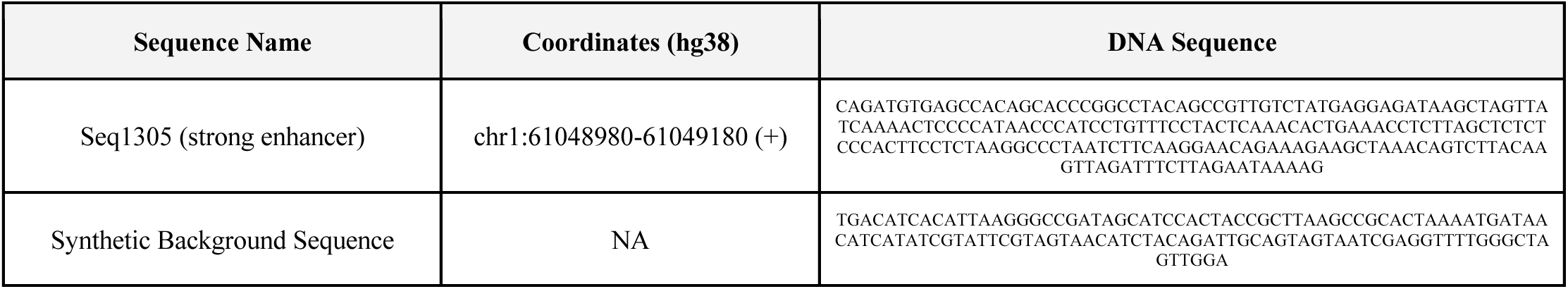
Information for the positive control and synthetic background DNA sequence used in the STARR-seq experiment. The positive control DNA sequence was previously validated in K562s using a Lenti-MPRA experiment^30^. The sequence was chosen due to a high activity score in K562, indicative of increased gene transcription. We also created a synthetic DNA sequence *in silico* to use as a background DNA sequence. The sequence was designed to have no TFBS or novel motifs in the sequence (see methods for details).

**Supplemental Figure S8.**
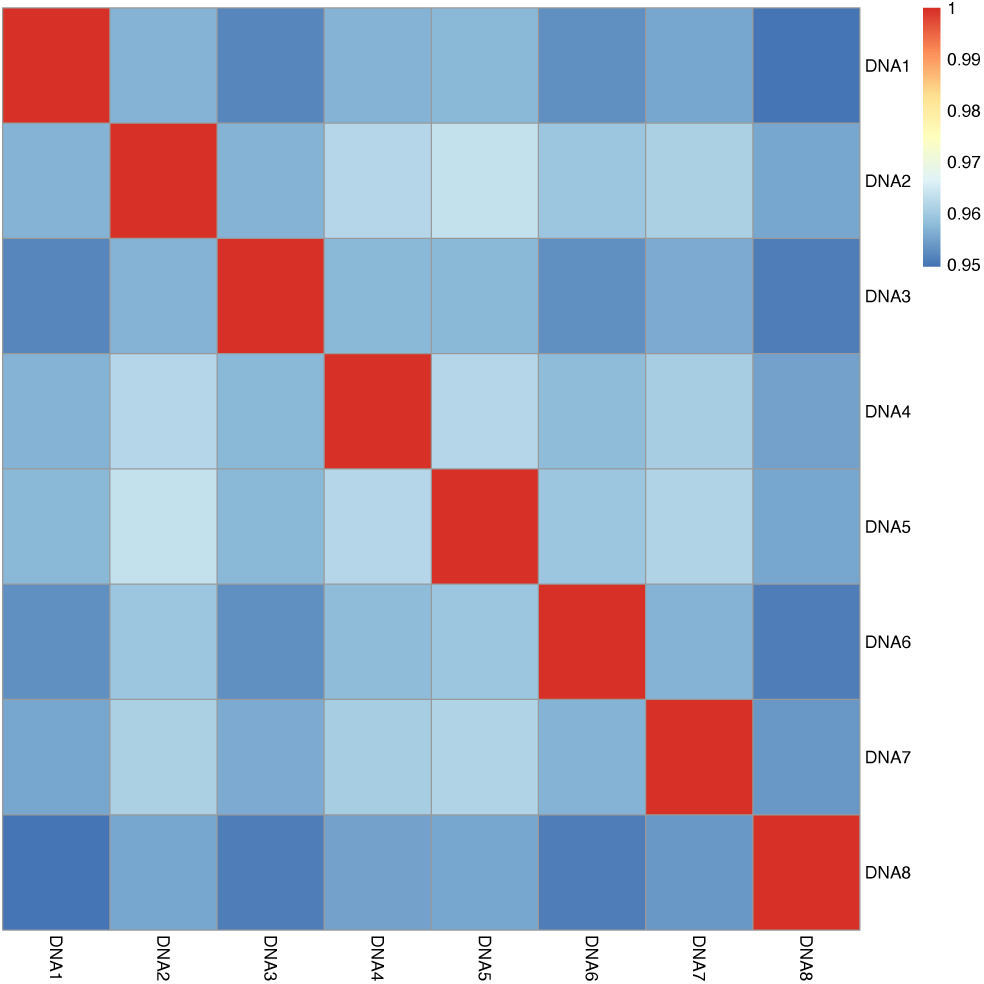
Pearson correlation (r) between sequencing replicates (n=8) obtained from sequencing the STARR-seq plasmid library.

**Supplemental Figure S9.**
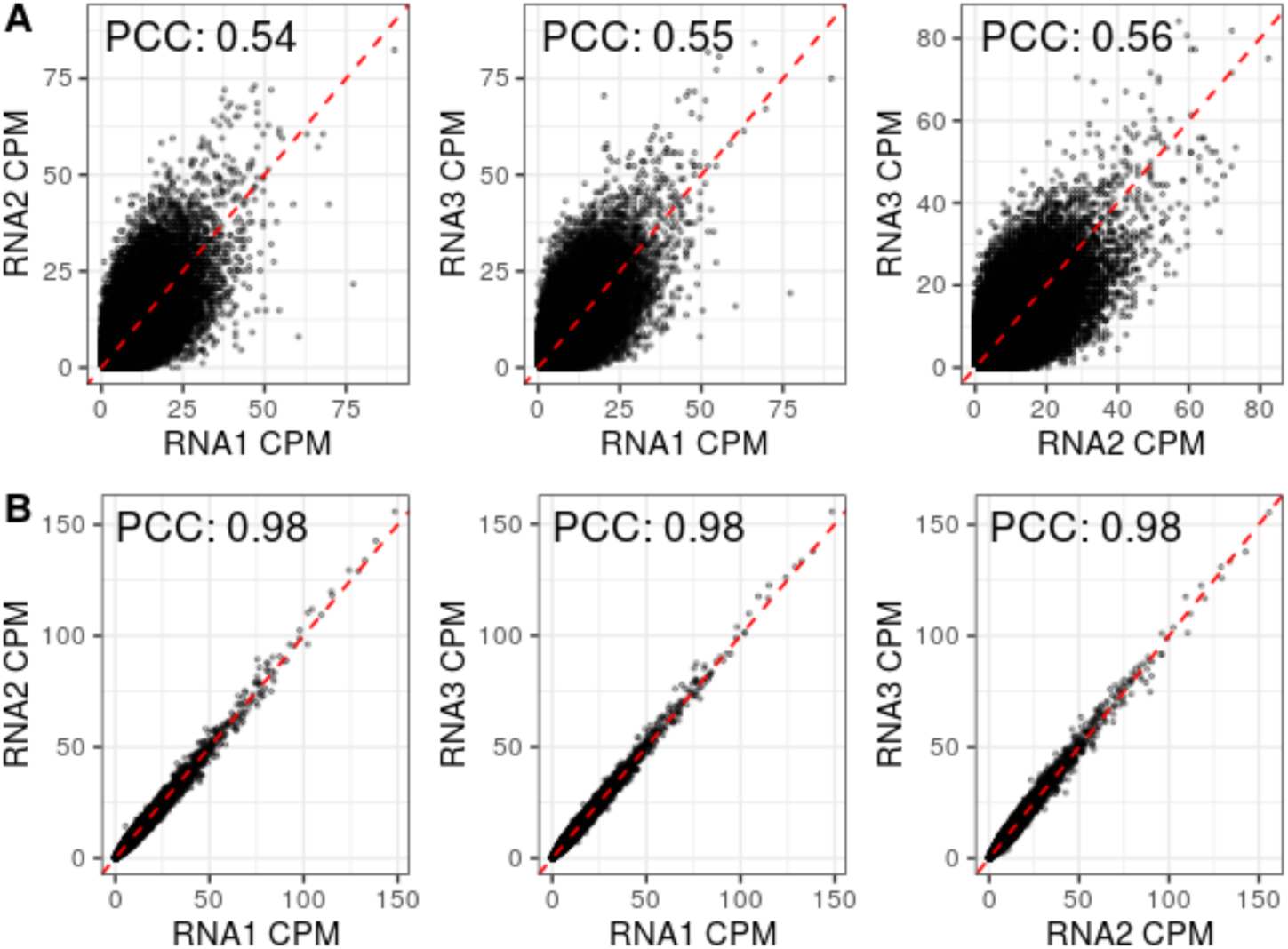
Pearson correlations (r) for counts per million (CPM) derived from amplicon-sequencing replicates (n=3) for transfected A) Jurkat and B) K562 cell lines.

**Supplemental Figure S10.**
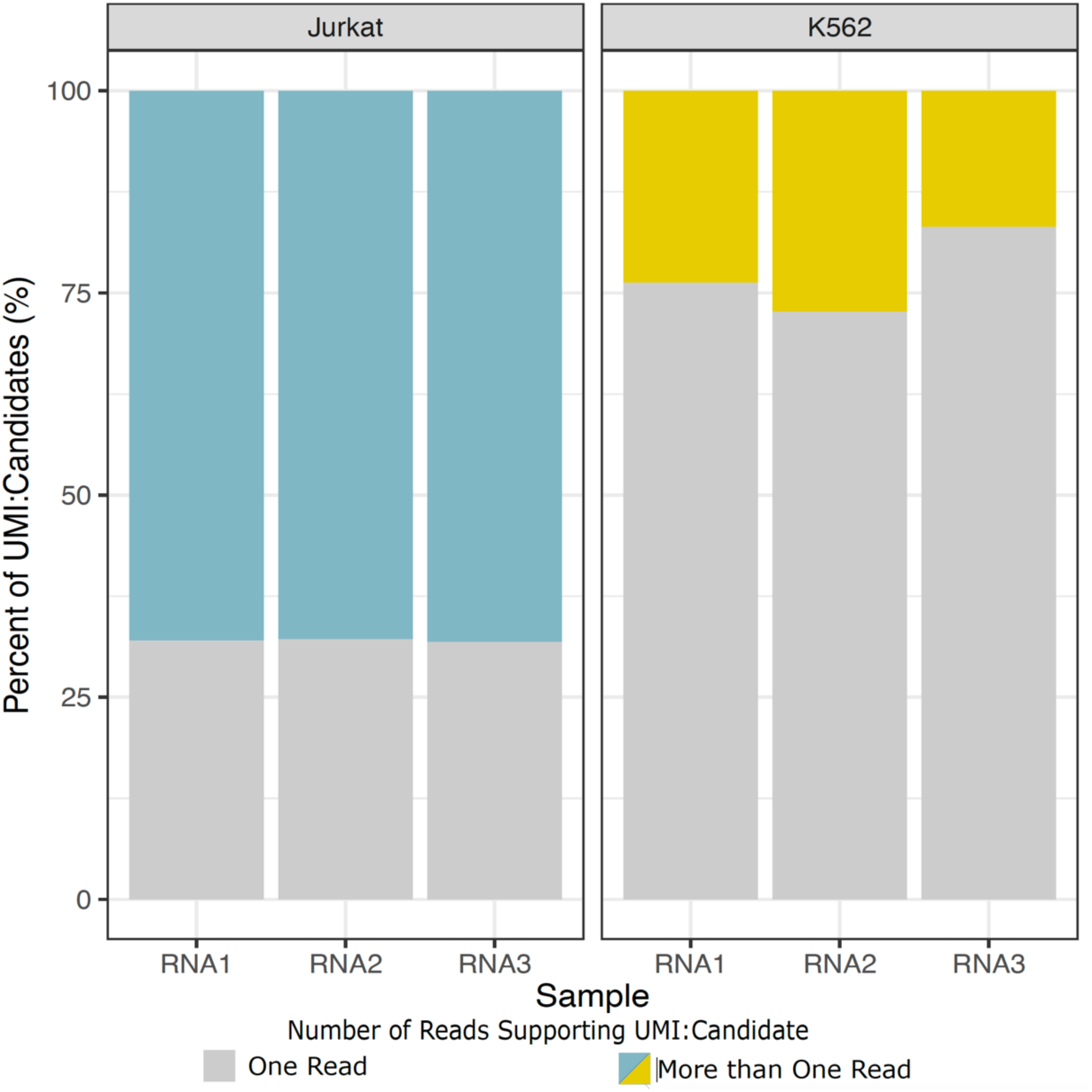
The plot shows the number of distinct oligo-UMIs that are covered by more than one read in Jurkat and K562 cells across the three replicates. Jurkats had sufficient coverage, whereas K562 cells could have used additional amplicon sequencing.

**Supplemental Figure S11.**
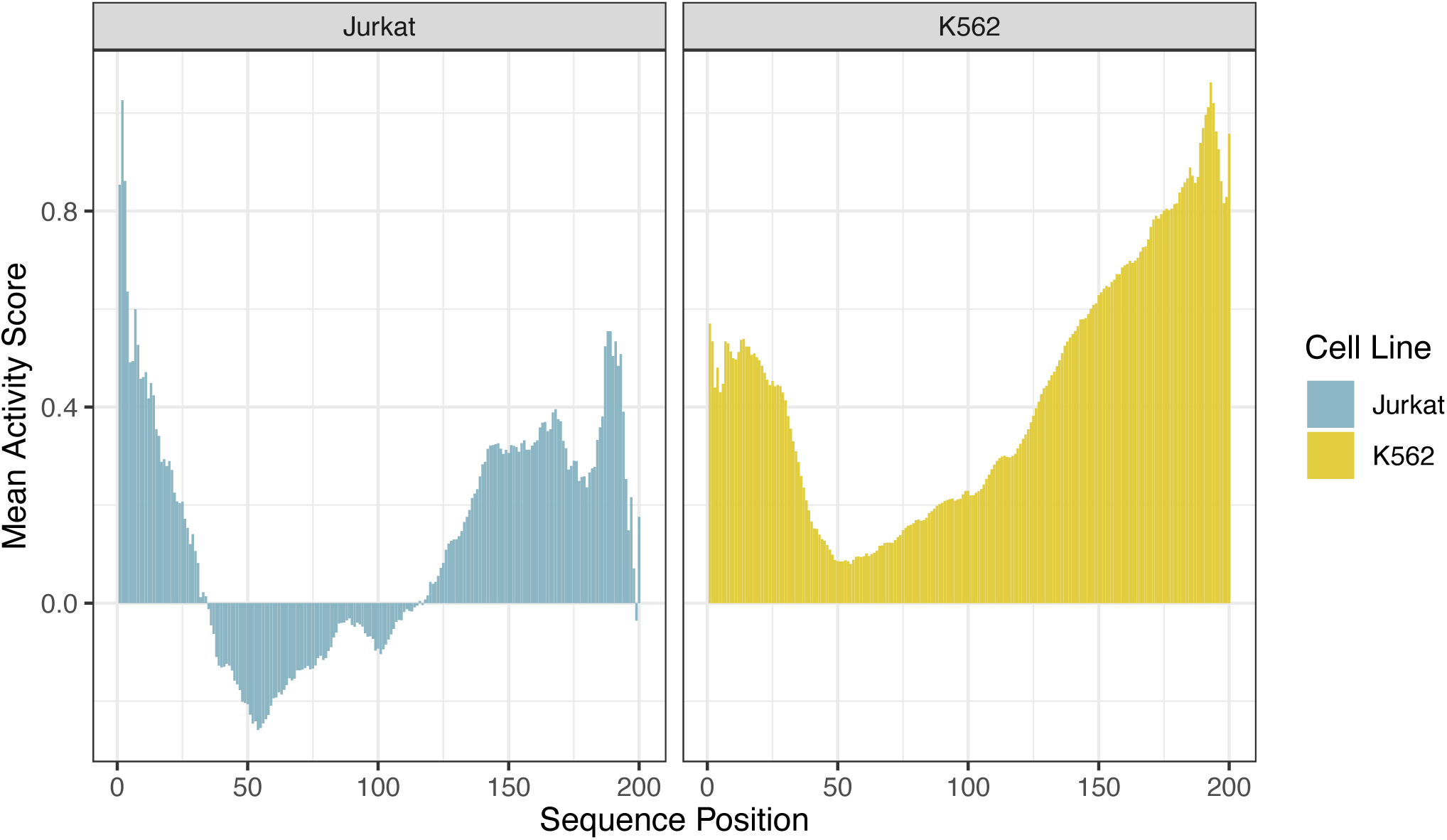
The positive control sequence (seq1305) drives transcription in Jurkat cells (left), as well as K562 cells (right). Most of the activity is driven by the 3’ end of the enhancer sequence.

**Supplemental Figure S12.**
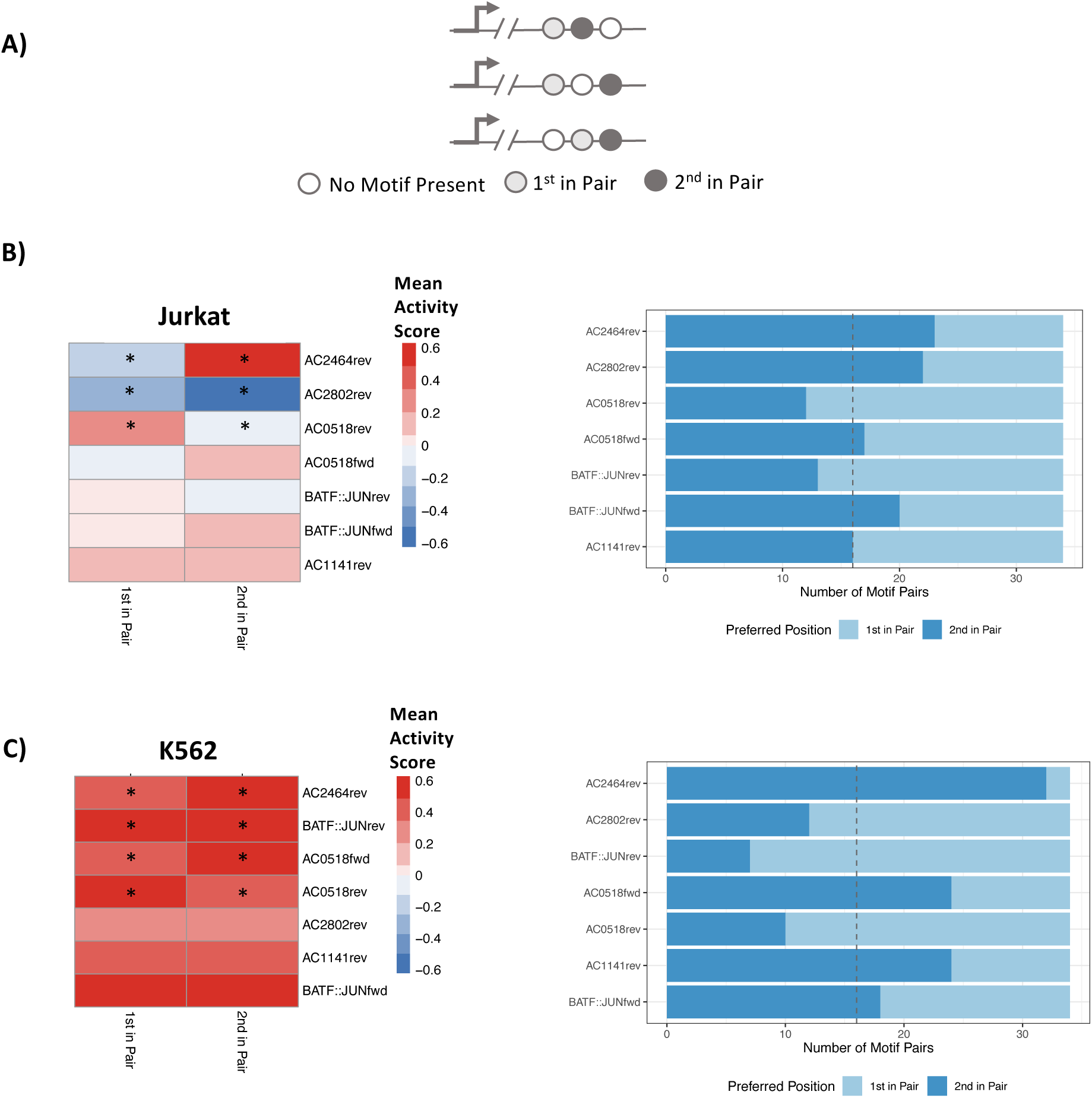
The position of a motif relative to the core-promoter has a significant effect on gene transcription. A) Schematic of the oligonucleotides used for the analysis. Only oligos containing a single copy of two distinct motifs were used to determine if the motifs of interest have a different effect on gene transcription depending on whether they are the first or second motif in the oligo relative to the core-promoter. B) The position of the motif relative to the core-promoter has a significant effect on gene transcription (* Wilcoxon t-test, Bonferroni corrected, p-value < 0.007) in B) Jurkat and C) K562 cells, left. The preference for the position is displayed in the figure on the right for both Jurkat and K562 cells.

**Supplemental Figure S13.**
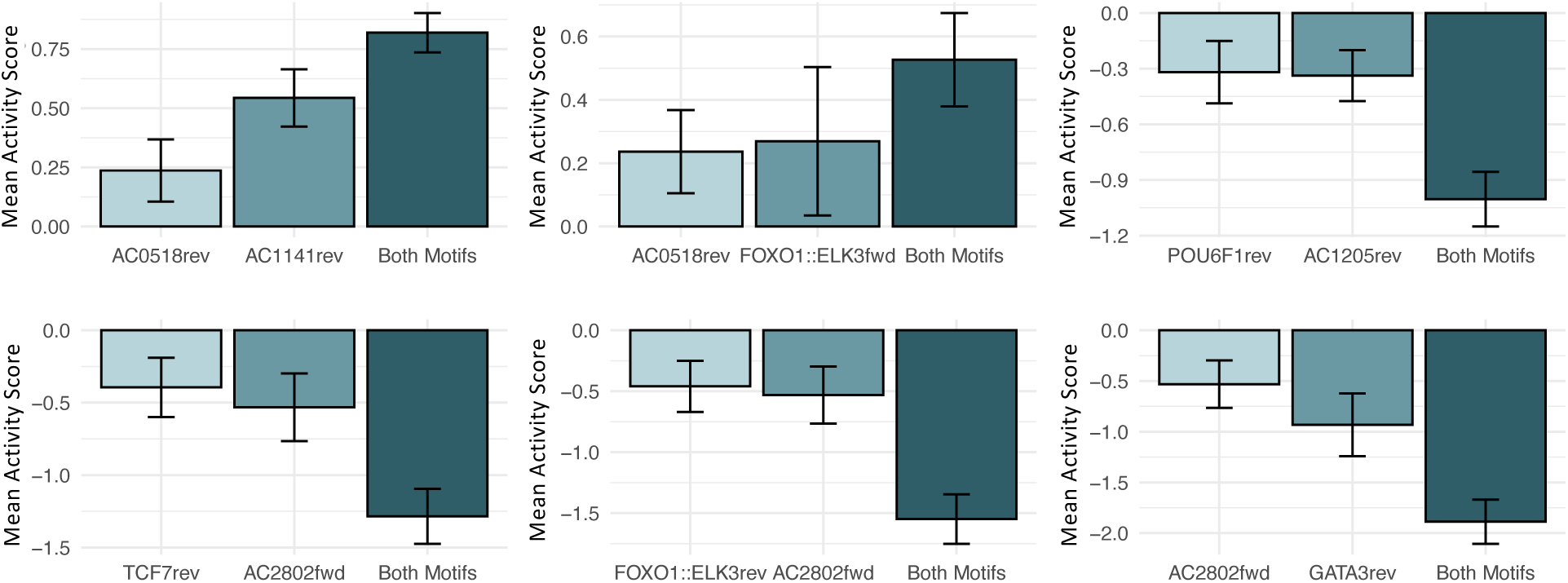
Pairs of motifs have additive effects on gene transcription. The activity scores for the pairs of motifs that have additive effects on gene transcription in Jurkats. Error bars represent the standard error.

**Supplemental Table S3.**
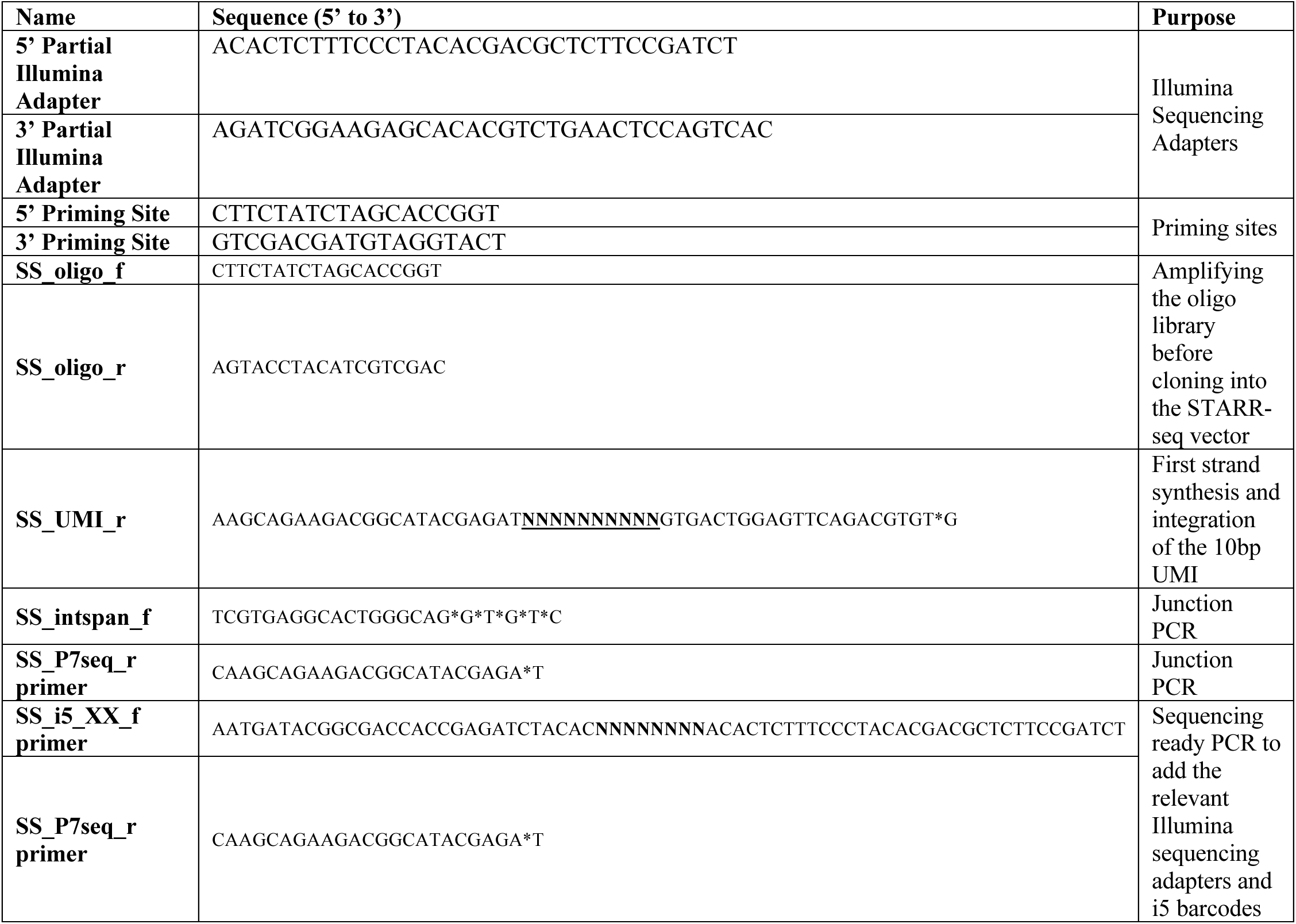
List of adapter and primer sequences used in the STARR-seq assay. * = phosphorothioate bond to protect against 3’-5’ exonuclease activity. **<u>NNNNNNNNNN</u>** = unique molecular identifier. **NNNNNNNN** = i5 barcodes. XX corresponds to the two-digit number corresponding to the i5/i7 primer pair UDI ID from Illumina (UDI00XX), where the i5 barcodes were extracted from.

**Supplemental Table S4.** Sequencing statistics for the DNA and amplicon libraries.

| **Sample** | **Number of Paired-End Reads** | **Average Number of Paired-End Reads Per Oligo**<br>**(n=59,776)** |
| --- | --- | --- |
| **DNA1** | 20,176,782 | 337 |
| **DNA2** | 27,200,642 | 455 |
| **DNA3** | 20,914,051 | 349 |
| **DNA4** | 25,458,490 | 425 |
| **DNA5** | 26,854,834 | 449 |
| **DNA6** | 21,413,241 | 358 |
| **DNA7** | 23,746,091 | 397 |
| **DNA8** | 18,430,233 | 308 |
| **Jurkat RNA1** | 4,362,984 | 72 |
| **Jurkat RNA2** | 3,479,663 | 58 |
| **Jurkat RNA3** | 3,153,281 | 52 |
| **K562 RNA1** | 11,631,784 | 194 |
| **K562 RNA2** | 10,436,332 | 174 |
| **K562 RNA3** | 9,776,950 | 163 |

